# Gene Expression Correlates of the Cortical Network Underlying Sentence Processing

**DOI:** 10.1101/439984

**Authors:** Xiang-Zhen Kong, Nathalie Tzourio-Mazoyer, Marc Joliot, Evelina Fedorenko, Jia Liu, Simon E. Fisher, Clyde Francks

## Abstract

A pivotal question in modern neuroscience is which genes regulate brain circuits that underlie cognitive functions. However, the field is still in its infancy. Here we report an integrated investigation of the high-level language network (i.e., sentence processing network) in the human cerebral cortex, combining regional gene expression profiles, task fMRI, large-scale neuroimaging meta-analysis, and resting-state functional network approaches. We revealed reliable gene expression-functional network correlations using three different network definition strategies, and identified a consensus set of genes related to connectivity within the sentence-processing network. The genes involved showed enrichment for neural development and actin-related functions, as well as association signals with autism, which can involve disrupted language functioning. Our findings help elucidate the molecular basis of the brain’s infrastructure for language. The integrative approach described here will be useful to study other complex cognitive traits.

## Introduction

A pivotal question in modern neuroscience is which genes regulate brain circuits that underlie cognitive functions. In the past decade, imaging genetics has provided a powerful approach for exploring this question in humans, by combining neuroimaging data and genotype information from the same subjects and searching for associations between interindividual variability in neuroimaging phenotypes and genotypes within a sample. While numerous imaging genetics studies have now been published (Avinun, Nevo, Knodt, Elliott, & Hariri, 2018; Bigos & Weinberger, 2010; Carter et al., 2017; Thompson et al., 2014), there remain key issues which affect the field, including sample size limitations, the need to correct for multiple comparisons, and the small effect sizes that are typical of associations with common gene variants.

Recently, researchers have begun to probe the molecular genetic architecture of the human brain not only through genotypes and inter-individual variability, but also using regional gene expression mapping in post mortem brain tissues, in combination with neuroimaging data collected from living individuals (Bartres-Faz et al., 2019; Kong, Song, Zhen, & Liu, 2017; McColgan et al., 2018; Richiardi et al., 2015; Romero-Garcia, Warrier, Bullmore, Baron-Cohen, & Bethlehem, 2018; Romme, de Reus, Ophoff, Kahn, & van den Heuvel, 2017; Vertes et al., 2016). This approach has brought important new advances: transcriptional profiles have been linked to neural architecture with respect to both functional connectivity measured during the resting state (also called intrinsic connectivity) (Richiardi et al., 2015; Vertes et al., 2016) and structural connectivity (Fulcher & Fornito, 2016; Richiardi et al., 2015), as well as to alterations of connectivity in brain disorders such as schizophrenia (Romme et al., 2017), autism spectrum disorder (ASD) (Romero-Garcia et al., 2018), and Huntington’s disease (McColgan et al., 2018). A necessary assumption is that the group-averaged gene expression map, within a given post mortem dataset, is sufficiently representative to be matched with the group-averaged neuroimaging-derived map from a set of living subjects.

The present study was inspired by a number of recent studies that probed connectivity data in relation to inter-regional gene expression similarity. Richiardi and colleagues (2015) used data from resting-state functional magnetic resonance imaging (rs-fMRI) to show that the network modularity structure, derived from functional connectivity patterns across the cortex, was correlated with inter-regional similarity of gene expression profiles. Specifically, regions within a module (i.e., regions with stronger functional connectivity) showed more similar gene expression profiles than across different modules. French & Pavlidis (2011) showed that average gene expression similarity is higher for regions with structural connections, compared to unconnected regions, in adult rodent brain (French & Pavlidis, 2011), while Goel et al. (2014) examined the association between gene expression similarity and pairwise structural connectivity in the human brain (Goel, Kuceyeski, LoCastro, & Raj, 2014). Seidlitz et al. (2018) showed a significant correlation between inter-regional gene co-expression and edge weights based on morphometric similarity, in the human brain (Seidlitz et al., 2018). In another study, Romme et al. (2016) investigated the transcriptional profiles of a set of genes known to contain inherited variants associated with schizophrenia, and found that they were significantly correlated with regional reductions in the strength of white matter connections in people with schizophrenia. Whitaker et al. (2016) investigated links between transcriptome data and adolescent cortical development, specifically cortical thickness shrinkage and intracortical myelination, and implicated oligodendroglial genes in these processes, as well as a set of genes associated with risk for schizophrenia (Whitaker et al., 2016). More recently, gene expression in the human brain has also been linked to regional differences in cortical scaling (Reardon et al., 2018), and anatomical hierarchy (Burt et al., 2018), using similar spatial correlation approaches. For a current review on linking neuroimaging and transcriptome, see (Fornito, Arnatkeviciute, & Fulcher, 2019).

These previous studies combining gene expression and brain imaging data have provided strong evidence that patterns of gene expression co-vary with anatomical and functional organization of the human brain. However, the ultimate goal is to gain a rich and detailed picture of the genetic and molecular mechanisms that support each core cognitive ability. Here we will focus on the quintessential and human-unique capacity for language. Previous studies have suggested that language-related cognitive performance is highly heritable (e.g., (Dale et al., 1998; Guen, Amalric, Pinel, Pallier, & Frouin, 2018; Newbury, Bishop, & Monaco, 2005)), and that brain activations associated with semantic comprehension tasks are also heritable (Guen et al., 2018). Moreover, genetic factors also play a substantial role in susceptibility to language-related neurodevelopmental disorders such as childhood apraxia of speech (Eising et al., 2018), developmental language disorder (specific language impairment) and developmental dyslexia (Deriziotis & Fisher, 2017). Crucially, although a small number of genes – such as *FOXP2* (e.g., (Fisher & Scharff, 2009; Lai, Fisher, Hurst, Vargha-Khadem, & Monaco, 2001)) – have now been unambiguously linked to language-related disorders, these genes cannot by themselves explain the large majority of heritable variation, nor can they conceivably create or maintain the necessary brain circuits underlying language without interacting with a large number of other genes (Graham & Fisher, 2015; Konopka & Roberts, 2016).

In addition, linguistic deficits are often found with heritable, neurodevelopmental disorders for which impaired language function is not necessarily diagnostic, including intellectual disability, autism spectrum disorder (ASD), and schizophrenia (Bearden et al., 2000; Geschwind & Flint, 2015; Kaufman, Ayub, & Vincent, 2010; Kleinhans, Muller, Cohen, & Courchesne, 2008; Lombardo et al., 2015; Schizophrenia Working Group of the Psychiatric Genomics, 2014; Tager-Flusberg, Paul, & Lord, 2005; Tomblin, 2011). Linguistic ability also correlates with intelligence in the general population (Silva, Williams, & Mcgee, 1987). Thus, identifying genes and molecular mechanisms associated with language will not only i) yield a better understanding of the biological pathways that lead to the emergence of language phylo- and onto-genetically, but also ii) help identify susceptibility factors for language impairments in neuropsychiatric conditions, which could lead to improved diagnostic and treatment strategies.

To shed further light on the molecular architecture underpinning language circuits, here we synergistically combined task fMRI data, resting-state functional connectivity, and gene transcription profiles in the human brain. Specifically, we targeted sentence-level processing as an essential, high-level linguistic function, which has been linked to a network of regions particularly in the left temporal and frontal cortices (Dronkers, Wilkins, Van Valin, Redfern, & Jaeger, 2004; E. Fedorenko & Thompson-Schill, 2014; L. Labache et al., 2018; Price, 2012; Vigneau et al., 2006), as opposed to lower level language-related functions which can rely, for example, on primary auditory and motor areas. First, we defined the cortical regions for left hemispheric sentence processing based on task fMRI data, using three different sets of criteria and data to ensure robustness and generalizability across approaches. Then, we estimated the resting-state functional connectivity networks among these regions, using rs-fMRI data from two independent datasets. Previous studies have shown that functionally defined language-responsive regions form an integrated system also in the resting state (Blank, Kanwisher, & Fedorenko, 2014; Cole, Bassett, Power, Braver, & Petersen, 2014; L. Labache et al., 2018). Next, we examined the correlations between these functional connectivity patterns and the corresponding inter-regional similarity patterns of gene expression, based on post mortem data. Similar pairwise connectivity/similarity correlation approaches have been applied in a number of previous studies aiming to probe connectivity data in relation to gene expression data (see above). We then assessed the contributions of each individual gene to the observed correlations, and identified a consensus set of top genes (showing positive contribution scores) across all six analyses (i.e., three definition strategies for the sentence processing regions by two rs-fMRI datasets for estimating the functional connectivity). Finally, using several bioinformatics databases, we explored the biological roles and expression specificity of these genes, and tested whether they showed an enrichment for association signals with ASD, schizophrenia or intelligence, using genome-wide association study (GWAS) data. We also analyzed three other functional networks by way of comparison to these language-related networks, which were the spatial navigation network, fronto-parietal multiple demand network, and default mode network. Given left-hemisphere dominance of the language network and limited post mortem gene expression data for the right hemisphere (see below), we focused on the left hemisphere in the present study.

## Materials and Methods

Fig. 1 shows a schematic of our approach for measuring the correlation between functional connectivity and gene expression profiles within a given network of brain regions. This analysis pipeline consisted of a) defining sets of cortical regions using task activation data (directly or via meta-analysis), b) estimating the resting-state functional connectivity and gene expression similarity among these regions, and c) assessing the correlation between the functional connectivity and gene expression networks, along with subsequently d) estimating each gene’s individual contribution using a ‘leave-one-out’ approach (see below). See below for details of each dataset and procedure.

**Fig. 1.**
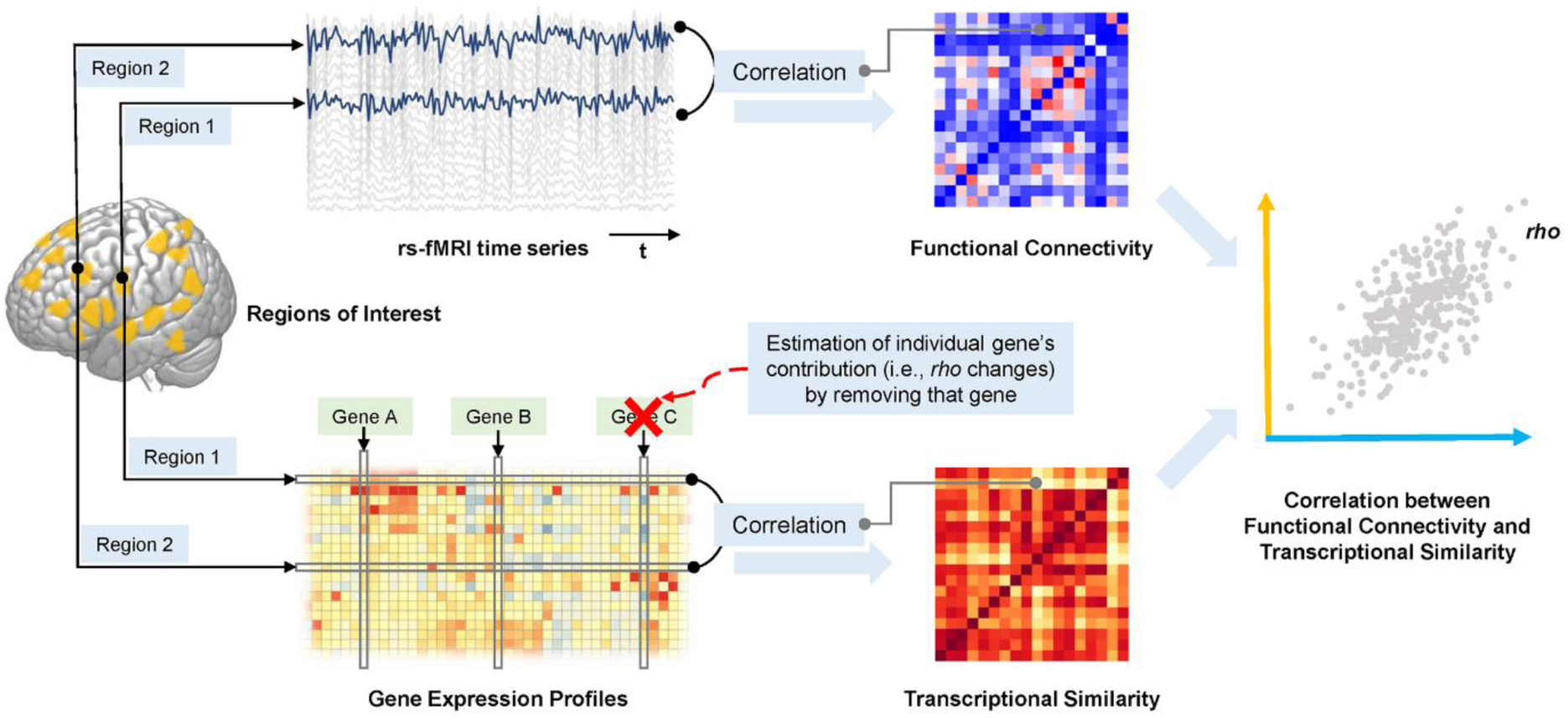
Schematic of the pipeline for computing the correlation between resting-state functional connectivity and transcriptomic similarity, within a network of regions first defined according to task fMRI data. Regional time series from rs-fMRI were used to estimate functional connectivity between each pair of regions (in the upper matrix, red indicates stronger functional connectivity; blue indicates weaker functional connectivity). Regional gene expression profiles were applied to estimate inter-regional transcriptional similarity (in the lower matrix, red indicates higher similarity; yellow indicates lower similarity; lower panel). Resting-state functional connectivity patterns (orange axis in the right-hand panel) are correlated with patterns of transcriptional similarity (blue axis in the right-hand panel). Each gene’s contribution to this correlation is estimated by removing that gene at the step of gene expression network construction (red cross) and repeating the subsequent analysis.

### Datasets

fMRI dataset acquisitions were approved by the Institutional Review Board of each site. Written informed consent was obtained when necessary from all participants (*N* = 244), before they took part.

#### BIL&GIN

This dataset included 144 healthy right-handed adults (aged 27±6 years; 72 females) drawn from the larger BIL&GIN database which is roughly balanced for handedness (Mazoyer et al., 2016). Each participant completed three slow-event fMRI runs (gradient echo planar imaging, TR = 2.0 s, acquisition voxel size = 3.75 × 3.75 × 3.75 mm^3^; 3T Philips Intera Achieva scanner) in which they were asked to complete 3 different sentence tasks including covertly producing, listening to, or reading sentences and familiar word lists as reference. Of these participants, 137 also completed rs-fMRI scans using the same imaging sequence as that used for the tasks, which lasted 8 minutes (240 volumes). Immediately prior to rs-fMRI scanning, the participants were instructed to “keep their eyes closed, to relax, to refrain from moving, to stay awake and to let their thoughts come and go”. Note that the latter scanning session took place around 1 year before the task fMRI scans. For more details see further below, and (Loic Labache et al., 2018; L. Labache et al., 2018).

#### NeuroSynth

Neurosynth (http://neurosynth.org) is a platform for large-scale synthesis of task fMRI data (Yarkoni, Poldrack, Nichols, Van Essen, & Wager, 2011). It uses text-mining techniques to detect frequently used terms as proxies for concepts of interest in the neuroimaging literature: terms that occur at a high frequency in a given study are associated with all activation coordinates in that publication, allowing for automated term-based meta-analysis. Despite the automaticity and potentially high noise resulting from the large-scale meta-analysis, this approach has been shown to be robust and meaningful (e.g., Helfinstein et al., 2014; Kong, Song, et al., 2017; Kong, Wang, et al., 2017; Yarkoni et al., 2011), due to the high number of studies included. We used database version 0.6 (current as of July 2018) which included 413,429 activation peaks reported in 11,406 studies (see below for the search terms employed).

#### EvLabN60

This dataset included statistical maps of the task fMRI contrast for passively reading sentences versus non-words from 60 participants (aged from 19 to 45; 41 females; all right-handed). This fMRI task (TR = 2.0 s, acquisition voxel size 2.1 × 2.1 × 4.0 mm^3^; 3 T Siemens Trio scanner) was designed to localize the sentence processing network (for details, see Fedorenko et al., 2010). The sentences > non-words contrast has been previously shown to reliably activate language-sensitive regions and to be robust to the materials, task, and modality of presentation (E. Fedorenko, Behr, & Kanwisher, 2011; E. Fedorenko, Hsieh, Nieto-Castanon, Whitfield-Gabrieli, & Kanwisher, 2010; Mahowald & Fedorenko, 2016; Scott, Gallee, & Fedorenko, 2017). In addition, the EvLabN60 dataset was used to define two other networks used as comparisons to sentence processing networks (below). A spatial working memory task was designed to localize the fronto-parietal multiple demand system (i.e., contrast Hard versus Easy) (Blank et al., 2014; E. Fedorenko, Duncan, & Kanwisher, 2013) and the default mode network (i.e., contrast Easy versus Hard) (Leech, Kamourieh, Beckmann, & Sharp, 2011; McKiernan, Kaufman, Kucera-Thompson, & Binder, 2003; Park, Polk, Hebrank, & Jenkins, 2010). Participants were instructed to keep track of four (Easy condition) or eight (Hard condition) sequentially presented locations in a 3×4 grid. In both conditions, participants performed a two-alternative forced-choice task at the end of each trial to indicate the set of locations they just saw. For more details, see Fedorenko et al., 2011.

#### GEB

GEB (http://www.brainactivityatlas.org), which is an abbreviation for “Gene-Environment-Brain-Behavior”, provided an independent rs-fMRI dataset for the present study. GEB is an on-going project that focuses on linking individual differences in human brain and behaviors, to environmental and genetic factors (Kong, Song, et al., 2017; Wang et al., 2016; Zhen et al., 2017; Zhen et al., 2015). Rs-fMRI data from forty college students (20 females; aged = 20.3 ± 0.91 years) were included in this study. The resting-state scan lasted 8 min and consisted of 240 contiguous echo-planar-imaging (EPI) volumes (TR = 2.0 s; acquisition voxel size = 3.125 × 3.125 × 3.6 mm^3^; 3 T Siemens Trio scanner). During the scan, participants were instructed to relax and remain still, with their eyes closed. The dataset has high quality in terms of minimal head motion and registration errors, and has been used in several previous studies (e.g., Kong, Wang, et al., 2017; Wang et al., 2016).

#### AHBA

AHBA (Allen Human Brain Atlas; http://www.brain-map.org) is a publicly available online resource for gene expression data. The atlas characterizes gene expression (i.e. messenger RNA quantification) in postmortem human brain based on genome-wide microarray-based measurement, for over 20,000 genes at ~500 sampling sites distributed over the whole brain. See (M. J. Hawrylycz et al., 2012) for more details about the data collection. Normalized expression data were used in the present study. To date (search conducted on Mar. 30, 2017), six adult donors with no history of neuropsychiatric or neurological conditions were available in the database (age 24, 31, 34, 49, 55, and 57 years; 1 female). Left hemisphere cerebral cortical data are available for all six donors whereas right-hemisphere data are available for only two of them. Detailed information on donors and analysis methods is available at www.brain-map.org. Structural brain imaging data of each donor were used to align sampling sites into standard coordinate space. We also used gene expression data from the Allen Institute’s BrainSpan project (http://www.brainspan.org/), which includes human brain tissues from age 8 weeks post conception to 40 years, sampling an average 13 regions (range, 1-17) from one to three brains per time point, and measured using RNA-sequencing (Li et al., 2018). We used these latter data for examining the expression of specific genes of interest across human brain development.

#### Genome-wide Association Scan results for ASD, Schizophrenia and intelligence

We downloaded GWAS summary statistics from the Psychiatric Genomics Consortium (http://www.med.unc.edu/pgc) for ASD with up to 7387 cases and 8567 controls (ASD GWAS 2017) (Autism Spectrum Disorders Working Group of The Psychiatric Genomics, 2017), and schizophrenia with up to 36,989 cases and 113,075 controls (PGC-SCZ2) (Schizophrenia Working Group of the Psychiatric Genomics, 2014). We also downloaded GWAS association results for intelligence, based on 78,308 individuals from the UK Biobank, CHIC consortium, and five additional cohorts (https://ctg.cncr.nl/software/summary_statistics) (Sniekers et al., 2017). This kind of data comprises summary statistics from genetic association testing between each of millions of single nucleotide polymorphisms distributed over the genome, in relation to the trait or disorder of interest.

### Defining Cortical Regions for Sentence Processing

We employed three strategies and data sources for defining the cortical sentence processing network. To refer to the regions defined under each of these three approaches, we will use the terms Supramodal Sentence Areas (SmSA), Synthesized Sentence Areas (SSA), and One-contrast Sentence Areas (OcSA). Given left-hemisphere dominance of language network (see Introduction) and limited post mortem gene expression data for the right hemisphere (see above), we focused on the left hemisphere in the present study. Note that we made use of existing, previously published task-fMRI datasets for this study (see below), and do not repeat all of the method details here.

#### SmSA

We applied the definition of left-hemispheric high-order and supramodal sentence areas provided by The SENSAAS atlas (L. Labache et al., 2018). This atlas of language integrative and supramodal areas, involving 142 healthy rigth-handers, is based on the conjunction of activation across sentence production, listening and reading, as contrasted with activation for lists of overlearned words (again presented as either production, listening or reading tasks) in the same participants. Then, a second criterion was applied whereby leftward activation asymmetry was required during the 3 sentence minus words contrasts. See Labache et al. (2018) for a full description of this definition approach. Task fMRI data were used from the BIL&GIN dataset (see above) for this purpose. In order to obtain accurate measures of functional asymmetry, this work is based on the use of the AICHA atlas, including left-right homotopic regions of interest based on resting-state functional connectivity data (Joliot et al., 2015). In the 179 left-right pairs of homotopic regions of the AICHA atlas, fMRI signal variation and asymmetry were calculated for each task contrast and each participant. Regions with both significant activation and leftward asymmetry across the three sentence-level versus word-list task contrasts were identified. A significance threshold of Bonferroni-corrected *p* < 0.05 was applied. Thirty-two left-hemisphere regions were obtained, including 27 cortical and 5 subcortical regions (L. Labache et al., 2018).

**SSA** were defined based on a large-scale neuroimaging meta-analysis of fMRI studies using Neurosynth (see above). A combination of terms related to sentence processing were used, including “sentence comprehension”, “sentence”, and “sentences” (411 studies). The resulting meta-analysis map (i.e., the likelihood map that shows there would be activation in some specific brain regions given the presence of particular terms) was used in this study to cover regions that are relevant to the network of interest. To control the false positive rate in the statistical map, a false discovery rate (FDR) threshold was used of 0.01 on a whole brain basis. As for SmSA, the AICHA atlas was used to define the areas for functional network construction. Specifically, if a region from the AICHA atlas had more than half (50%) of its voxels showing significant specificity based on the thresholded mask from the meta-analysis (FDR-corrected *p* <0.01), we included that region as one of the SSA. The rationale for using the AICHA atlas was twofold. First, the AICHA atlas was generated using rs-fMRI connectivity data, with each region showing homogeneity of functional temporal activity within itself. Second, we wanted to ensure that the results were directly comparable with the results using SmSA.

**OcSA** were defined based on the probabilistic activation map of a single fMRI contrast from the EvLabN60 dataset (see above). The passively reading task has been previously shown to reliably activate language-sensitive regions and to be robust to the materials, task, and modality of presentation (E. Fedorenko et al., 2011; E. Fedorenko et al., 2010; Mahowald & Fedorenko, 2016; Scott et al., 2017). The map was created by overlapping the statistical maps from all participants for the contrast of sentences versus non-words (t >2.3) onto the MNI152 template, and then dividing by the total number of participants (e.g., (Zhen et al., 2017)). The value for each voxel in the obtained map indicated the probability of the voxel showing a significant contrast activation across the population. A probability threshold of 50% was applied to identify voxels showing consistent activation (t > 2.3) across subjects, and regions from the AICHA atlas with more than half of the voxels activated were included, as for SSA.

#### Exclusion of regions

We excluded one region which was relatively small (less than 150 voxels; i.e., G_Cingulum_Post-3) and had limited gene expression data in the AHBA (fewer than 2 sampling sites; see below), as well as subcortical areas which are known to have very different gene expression profiles to cerebral cortex and would swamp the analysis (e.g., Hippocampus), a region that had resting-state data missing in the GEB dataset (i.e., G_Paracentral_Lobule-4), and two deep regions where the gene expression data was found to diverge substantially from most cerebral cortical regions (i.e., G_ParaHippocampal-1 and G_Insula-anterior-1) (Fig. S1). This resulted in 21 SmSA, 22 SSA, and 12 OcSA. The same criteria were also applied for defining the comparison systems below (Table S1).

#### Comparison networks

The NeuroSynth term ‘navigation’ was used to localize cortical regions involved in spatial navigation (64 studies; search conducted on Nov. 3, 2016). A FDR threshold of 0.01 was used to control the false positive rate. Moreover, the default mode network and the fronto-parietal multiple demand networks were defined using the EvLabN60 dataset (see above), with the probabilistic activation maps of fMRI contrast Easy versus Hard, and Hard versus Easy, respectively, based on the spatial working memory task. Again, a probability threshold of 50% was applied to identify voxels showing consistent activation (t > 2.3) across subjects. Next, the AICHA atlas was used to identify regions for each functional network. Our purpose was to define a similar number of top regions for each comparison network, to support similarly-powered analyses of all networks in the downstream analyses. In order to obtain comparable numbers of top regions for each network, different thresholds of overlap were applied for each network. Specifically, we found that a threshold of 1/4 defined 19 top regions of the spatial navigation network (SNN) (see below), a threshold of 1/2 defined 17 top regions of the multiple demand network (MDN), and a threshold of 3/4 defined 12 top regions of the default mode network (DMN).

### Construction of the Functional Connectivity Networks

Two independent rs-fMRI datasets, i.e. BIL&GIN and GEB (above), were used for functional connectivity network construction among the sets of regions defined based on task fMRI activation.

#### Functional connectivity in the BIL&GIN dataset

The preprocessing of the BIL&GIN dataset was done by the Bordeaux group (MJ). Preprocessing procedures included head motion correction, registration onto the anatomical T1 image, the latter being stereotaxic registered on the MNI152 standard space. Additionally, time series for white matter and cerebrospinal fluid, the six head motion parameters, and the temporal linear trend were removed from the stereotaxic normalized rs-fMRI data using regression analysis and time series data were temporally filtered using a least squares linear-phase finite impulse response (FIR) filter design bandpass (0.01-0.1 Hz). For each participant and each region, a time series was then calculated by averaging the rs-fMRI time series of all voxels located within that region. For each individual, we computed the Pearson’s correlation coefficient between the time series of each pair of cortical regions from a given task-defined set of regions. Correlation coefficients were transformed to Gaussian-distributed z scores via Fisher’s transformation. For each of the 6 networks (3 sentence processing and 3 other functional networks), the functional connectivity matrix was computed by averaging data from each individual. Negative edges were set to zero before follow-up analysis. For a full description of the processing see (Doucet et al., 2011).

#### Functional connectivity in the GEB dataset

The preprocessing of the GEB dataset was done by the Beijing group (JL). Preprocessing procedures included head motion correction, spatial smoothing, intensity normalization, and removal of linear trend, using the FEAT preprocessing workflow implemented with Nipype (Gorgolewski et al., 2011). A temporal band-pass filter (0.01-0.1 Hz) was applied to reduce low frequency drifts and high-frequency noise. To eliminate physiological noise, such as fluctuations caused by motion or cardiac and respiratory cycles, nuisance signals were regressed out. Nuisance regressors were averaged cerebrospinal fluid signal, averaged white matter signal, global signal averaged across the whole brain, six head realignment parameters obtained by rigid-body head motion correction, and the derivatives of each of these signals. The 4-D residual time series obtained after removing the nuisance covariates were registered to MNI152 standard space. After preprocessing, a continuous time course for each region of a given task-based functional network was extracted by averaging the time courses of all voxels within that region. Temporal correlation coefficients between the extracted time course from a given regions and those from other regions were calculated to determine the strength of the connections between each pair of regions of a given functional network at rest. Correlation coefficients were transformed to Gaussian-distributed z scores via Fisher’s transformation to improve normality, resulting in a symmetric Z value matrix (i.e., functional connectivity) for each task-defined system of each participant. Due to the ambiguous biological explanation of negative correlations (Schwarz & McGonigle, 2011), we restricted our analyses to positive edges and set negative edges to 0. We have applied the same processing procedure in several previous studies (e.g., Kong, Wang, et al., 2017; Wang et al., 2016). After resting-state functional connectivity networks were obtained corresponding to each task-defined set of regions, a mean functional connectivity network for each set of regions was calculated by averaging across participants, which was then used for subsequent analyses.

### Construction of the Gene Expression Networks

Transcriptomic networks were constructed based on gene expression profiles in the human brain from the AHBA (above). Specifically, we first extracted the normalized expression score for each gene, from each sampling site and each donor. For genes for which the expression had been measured using multiple microarray probes in the AHBA data, average values were calculated per gene at each sampling site and in each donor. Given that AICHA atlas regions are defined in the standard MNI space, the location of each sampling site was then translated into the standard space using the *alleninfo* (https://github.com/chrisfilo/alleninf). Gene expression data from within a given region were then averaged per gene and across donors, to obtain a single expression measure of each gene per region. We restricted our analyses to cerebral cortical regions with at least two sampling sites summed across all donors, and data quality was then assured by only including genes showing relatively high inter-individual consistency of expression levels, i.e. we restricted our analyses to the top 5% of all genes based on differential stability across donors as assessed over the entire cerebral cortex (Fig. S2). This was a set of 867 genes that had differential stability greater than 0.357, as previously calculated (M. Hawrylycz et al., 2015). The concept of differential stability was quantified as the averaged Pearson correlation between pairs of brains over a set of cerebral cortical regions (M. Hawrylycz et al., 2015). The percentage threshold applied (5%) seems to be strict, but the expression consistency of most genes is relatively low in cortical regions in the Allen data (M. Hawrylycz et al., 2015), so that a 5% threshold still includes genes with stability across donors of less than 0.4 (Fig. S2). Excluding genes with very low inter-individual consistency was necessary for the purpose of linking average gene expression data from the Allen brain dataset to functional connectivity in other datasets. We also repeated our analysis using a lower stability threshold (i.e., top 10%, stability larger than 0.25; *N* = 1735 genes), in order to assess the robustness of our findings with respect to this threshold. As a negative control, we additionally repeated the analyses using the lowest 5% and lowest 10% of genes as regards differential stability across donors, with which we would expect null results. We also repeated the analyses based on random sampling (repeated 1000 times) of genes from the whole distribution of stability scores.

Our processing of the genetic data produced a vector of gene expression values across regions. Transcriptional similarity between pairs of regions was then estimated by Pearson correlation, as a measure of ‘transcriptomic connectivity’.

### Correlation between the Functional Networks and Gene Expression Networks

#### Network similarity analysis

We examined the correlations between functional connectivity networks and gene expression networks. The essential concept is that pairs of regions of a functionally specific network that have relatively higher similarity of their gene expression profiles could also show higher functional connectivity (see Introduction). Specifically, a vector was extracted from the upper triangle of the connectivity matrix of each network, and Spearman correlation was then calculated between the vectors of the functional connectivity networks and their corresponding gene expression networks. Unlike previous studies focusing on general organization of the human brain, e.g., modularity (Richiardi et al., 2015) and hubs (Vertes et al., 2016), the present approach tested direct correlation between functional connectivity and gene expression similarity within the functional networks of interest. We also calculated the partial correlations after controlling for the spatial distance between centers of regions (i.e., the Euclidean distance of MNI coordinates) within the AICHA atlas (Joliot et al., 2015).

#### Gene contribution index (GCI)

In addition to the overall correlations between functional connectivity and corresponding gene expression networks, we formulated a novel index, the gene contribution index (GCI), for estimating the contribution of each individual gene to an observed overall correlation. GCI was defined as the difference in the overall correlation before and after removing that gene at the step of gene expression network construction, i.e. based on a ‘leave-one-out’ approach. Note that this ‘leave-one-out’ procedure was applied at the step of gene expression network construction, rather than when performing correlation analysis between two matrices (Fig. 1). A similar approach has been applied in a recent study for the same purpose (i.e., estimation each gene’s contribution to the overall correlation) (Seidlitz et al., 2018).

#### Identification of ‘consensus genes’ correlated with the sentence processing network

We identified a set of ‘consensus genes’ (*N* = 41), i.e. those which had positive CGI scores in all six analyses of the sentence processing network, i.e. the three definition strategies SmSA, SSA, and OcSA, by the two independent rs-fMRI datasets BIL&GIN and GEB. These were therefore the genes which made the most consistent individual contributions to the significant overall correlations between expression similarity and functional connectivity, regardless of the definition of sentence network regions. For each of the three comparison networks (spatial navigation, fronto-parietal multiple demand, and default mode networks) we also identified the same number of genes (i.e., 41) showing the highest GCIs, based on the averaged score of each gene in the two analyses relevant to each of those networks, i.e., with the two rs-fMRI datasets.

### Follow-up Analyses of the Consensus Genes

We further investigated the consensus genes correlated with the sentence processing network by use of literature searches and bioinformatics tools:

#### Gene ontology

The gene ontology provides a classification scheme for genes based on what is known with respect to their molecular functions, the biological processes that they are involved in, or cellular components that they encode (http://www.geneontology.org/). Gene ontology analyses were performed with the Bioconductor package gProfileR (https://biit.cs.ut.ee/gprofiler/), using ontologies from Ensembl release 91. Gene sets containing between 25-1000 genes were included. All known genes were used for determining the statistical domain size in the analysis. The default g:SCS method in the tool was used for multiple testing correction (corrected *p* < 0.05).

#### Gene expression across brain development

We queried the Allen Institute’s BrainSpan project data (see Datasets above) for the consensus genes, in relation to their developmental changes in expression from embryo to adult.

#### Gene expression specificity

For the 41 consensus genes correlated with connectivity in the sentence processing network, we contrasted their expression levels between those cortical regions that were assigned to the sentence processing network under any definition (i.e. all SmSA, SSA and OcSA) (*N* = 31), versus all regions outside the system that were included among the SNN, MDN or DMN (*N* = 34). More information about the lists of regions can be found in Table S1. The independent samples t-test (equal variances not assumed) was used to contrast expression levels. Since multiple comparisons were performed across 41 genes, a significance threshold of FDR-corrected *p* value of 0.05 was applied (Benjamini-Hochberg FDR).

In terms of cell-type specificity, the expression levels of each of the 41 consensus genes, that were correlated with connectivity within the sentence processing network, were queried in a published dataset based on mouse cortical data (as indexed as sequence reads per kilobase of exon per million reads mapped (FPKM) (Zhang et al., 2014). We also queried another single-cell gene expression database (Zeisel et al., 2015) to investigate the cell-type specificity of genes. Several recently published single-cell, or single-nucleus, gene expression datasets from human neocortex also became available while this study was in progress (Fan et al., 2018; Lake et al., 2018; Li et al., 2018; Zhong et al., 2018). We also queried these datasets.

### Enrichment Analysis using GWAS Summary Statistics for ASD, Schizophrenia and Intelligence

We tested the hypothesis that the consensus set of sentence processing network genes was enriched for genetic association signals with ASD, schizophrenia or intelligence, in the publicly available results from very large-scale genome-wide association studies, based on case-control or general population cohorts. (There were no large-scale GWAS results yet available for reading/language measures in the general population, or disorders such as dyslexia or language impairment which involve linguistic deficits.) Specifically, we ran gene set analyses using the MAGMA software (Version 1.06; http://ctg.cncr.nl/software/magma). MAGMA was run with default settings. As each gene contains multiple individual polymorphisms within it, gene-based association scores were derived using the SNP-wise mean model, which considers the sum of - log(p-values) as derived from GWAS analysis, for single nucleotide polymorphisms (SNPs) located within the transcribed region of a given gene (using NCBI 37.3 gene locations). MAGMA accounts for gene-size, number of SNPs in a gene, and linkage disequilibrium (LD) between SNPs when estimating gene-based association scores. LD between SNPs was based on the 1000Genomes phase 3 European ancestry samples. In this analysis, the score for a given gene therefore indicates how strongly genetic variation within, or in linkage disequilibrium with, that gene is associated with the trait of interest (i.e., ASD, schizophrenia, or intelligence). These GWAS-based gene scores were subsequently used to compute gene set enrichment within the 41 consensus genes associated with the sentence processing network. The enrichment analysis tests whether the genes in a given set have, on average, higher GWAS-based gene scores than the other genes in the genome. No cutoff was made on the GWAS-based gene scores, so that for all genes the degree of association with the trait of interest was taken into account. A significance threshold of FDR-corrected *p* value of 0.05 was applied to correct for multiple comparisons (i.e., ASD, schizophrenia, or intelligence). Similarly, we conducted exploratory gene set analyses with top genes for the other comparison functional networks, to compare with results for the sentence processing network.

## Results

### Functional Networks

Given the absence of a universally agreed upon protocol for localizing brain regions that support high-level language processing (E. Fedorenko & Thompson-Schill, 2014), we used three different definition strategies: i) Supramodal Sentence Areas (SmSA) based on the concordance of activation across three language fMRI tasks and leftward lateralization completed by 144 healthy right-handers (L. Labache et al., 2018), ii) Synthesized Sentence Areas (SSA) based on large-scale neuroimaging meta-analysis of fMRI studies (Yarkoni et al., 2011), and iii) One-contrast Sentence Areas (OcSA) based on the probabilistic activation map of a single language-task fMRI contrast (E. Fedorenko et al., 2010) (see Methods). Regions were defined according to the AICHA brain atlas, which is derived from rs-fMRI connectivity data, with each region showing homogeneity of functional temporal activity within itself (Joliot et al., 2015). The three resulting maps of the sentence processing network included 21, 22, and 12 regions respectively, which showed considerable overlap, and were consistent with previous studies, especially as regards core language regions such as the temporal and frontal regions (Fig. 2; Table S1) (E. Fedorenko & Thompson-Schill, 2014).

**Fig. 2.**
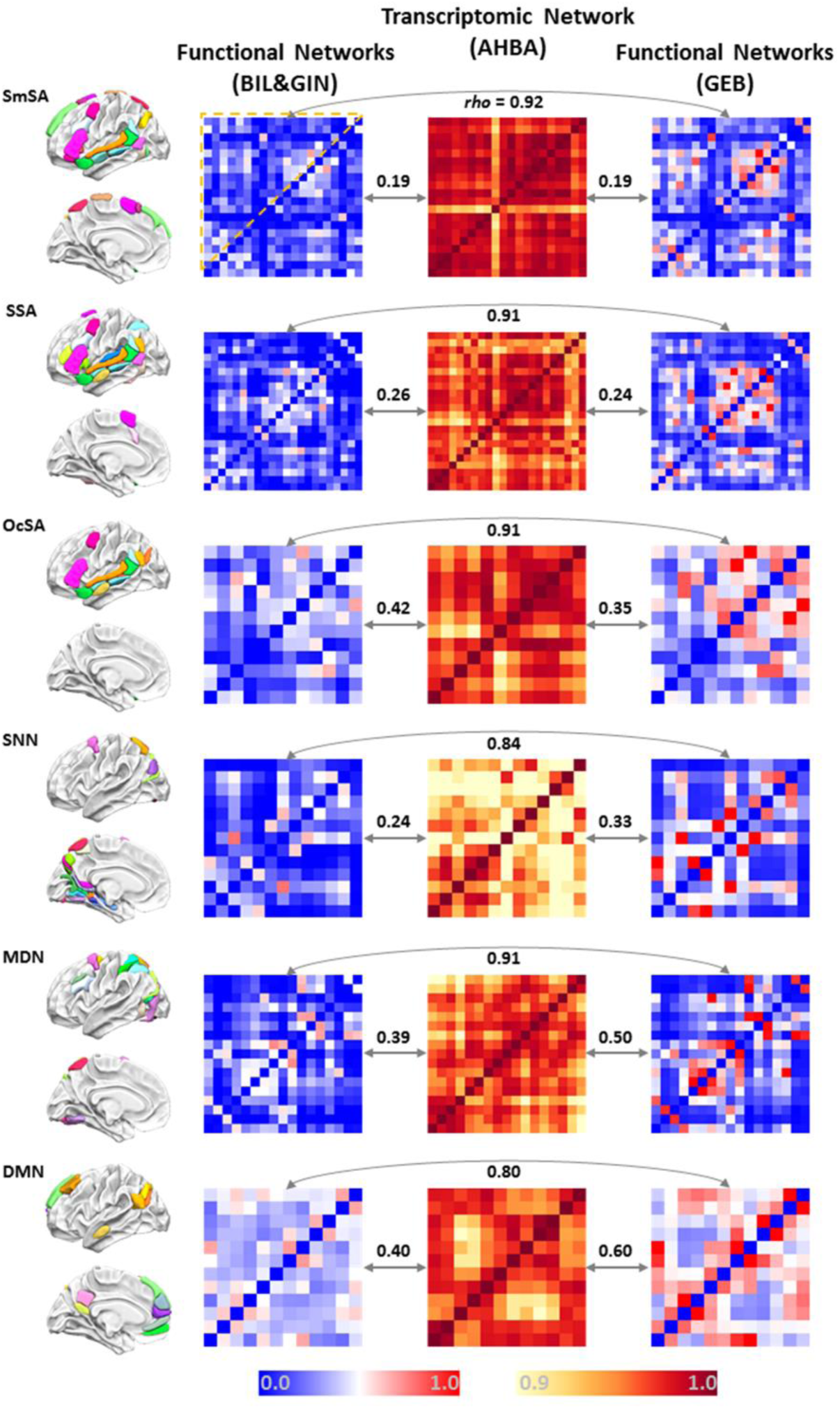
Functional and transcriptomic networks of the present study. First (left-side) column: the sets of cortical regions defined based on task fMRI data: top three (SmSA, SSA, and OcSA) correspond to the three different ways of defining the network for sentence processing, and bottom three correspond to the three other comparison networks. Second and fourth columns: the functional connectivity networks based on rs-fMRI from the BIL&GIN dataset and GEB dataset respectively. Third column: the transcriptomic networks based on the AHBA database. Within each matrix, the colors indicate the strength of pairwise inter-regional correlations or transcriptomic inter-regional similarity. Correlation values among the right three columns are the Spearman correlations (*rho*) of each pair of matrices for a given network, based on the upper triangle part of each matrix. All correlations were significant (*p* <0.01). SmSA, Supramodal Sentence Areas; SSA, Synthesized Sentence Areas; OcSA, One-contrast Sentence Areas; SNN, Spatial Navigation Network; MDN, Multiple Demands Network; DMN, Default Mode Network.

To compare to language-related networks, we also analyzed three other functional cortical networks: the spatial navigation network (SNN), fronto-parietal multiple demand network (MDN, similar to the dorsal attention network (Yeo et al., 2011)), and default mode network (DMN). The sets of cortical regions defined for these networks (see Methods) appeared consistent with previous literature (Buckner, Andrews-Hanna, & Schacter, 2008; Crittenden, Mitchell, & Duncan, 2016; Kong, Wang, et al., 2017) and showed little overlap with the sentence processing networks defined above (Fig. 2). We obtained 19, 17, and 12 areas for the SNN, MDN, and DMN respectively. More information about the spatial distribution of each definition and their overlaps can be seen in Table S1.

Connectivity within a given functionally-defined set of regions was estimated based on inter-regional synchronization of rs-fMRI time courses. The connectivity patterns based on two independent rs-fMRI datasets (BIL&GIN and GEB; see Methods) were highly correlated for all functional networks (*rho* > 0.80; Fig. 2), indicating high reproducibility of the group-level functional connectivity pattern with respect to AICHA atlas regions. Corresponding gene expression networks for each functionally defined set of regions were calculated based on post-mortem cortical gene expression data, and using pairwise similarities of regional gene expression profiles (Methods). To create a reliable average brain map of cortical gene expression, the analyses were restricted to 5% of all genes (i.e., 867 genes) which showed the highest differential stability of expression levels across individual donors, in cerebral cortical data from the Allen Brain Atlas (M. Hawrylycz et al., 2015). This approach meant that we focused only on genes which have consistent expression with respect to the average organization of the cerebral cortex. Within a given network, different pairs of regions varied in how similar they were in gene expression, although all regional pairwise correlations were high (greater than 0.9) (Fig. 2).

### Similarity between Functional Networks and Gene Expression Networks

We used correlation analysis to test whether regions with more similar gene transcription profiles show stronger resting-state functional connectivity, within each specific functional network. As expected, based on data from the BIL&GIN dataset, we found significant correlations between functional connectivity and the corresponding transcriptomic similarity patterns for each sentence processing network definition (Fig. 2; SmSA: *rho* = 0.19, *p* = 0.0048; SSA: *rho* = 0.26, *p* < 0.0001; OcSA: *rho* = 0.42, *p* = 0.00045). In addition, we obtained similar results for each of the comparison networks (Fig. 2; SNN: *rho* = 0.24, *p* = 0.0020; MDN: *rho* = 0.39, *p* < 0.0001; and DMN: *rho* = 0.40, *p* = 0.00081). Using connectivity data from the independent rs-fMRI dataset GEB, highly similar results were found (Fig. 2; SmSA: *rho* = 0.19, *p* = 0.0054; SSA: *rho* = 0.24, *p* = 0.00018; OcSA: *rho* = 0.35, *p* = 0.0042; SNN: *rho* = 0.33, *p* < 0.0001; MDN: *rho* = 0.50, *p* < 0.0001; and DMN: *rho* = 0.60, *p* < 0.0001). All gene-brain correlations survived correction for multiple testing (FDR corrected *p* < 0.01). We found similar correlations when applying a more inclusive threshold for gene expression differential stability across donors (stability > 0.25, i.e. the top 10% genes in stability), and as a negative control, we also tested the bottom genes in differential stability across donors (i.e., bottom 5% or 10% of genes), but saw little evidence of significant gene-brain correlation (Table S2), as expected. We also repeated the gene-brain correlation analyses based on random sampling (repeated 1000 times) of genes from the whole distribution of stability scores, and found that the correlations based on the top genes were greater than the correlations of random gene sets of equal size (Fig. S3), which further supported the observed gene-brain associations.

For the top 5% of genes by inter-donor stability, we also calculated the partial correlations between transcriptomic and functional connectivity matrices after correcting for the spatial proximity of regions (see Methods). These partial correlations were weaker than those without the correction for spatial proximity, and mostly non-significant (Table S3), suggesting that it is mostly local regional similarities that drive the overall transcriptomic-connectivity correlations within the functional networks defined in this study.

### Gene Contribution Index (GCI)

With a ‘leave-one-out’ procedure, we obtained ‘gene contribution index’ (GCI) scores for all individual genes, and for each functional network definition, which indicated the extent to which each gene affected the overall connectivity-transcriptome correlation for a given network (see Methods). To investigate the similarity of gene contribution patterns across different networks, we calculated the correlation between the GCI scores of each pair of networks. As expected, for the three different definitions of the sentence processing network, which involved largely overlapping sets of cortical regions, the GCI scores showed substantial correlations, which were also reproducible across the two independent rs-fMRI datasets (Fig. 3A; BIL&GIN: Mean *r* = 0.40, from 0.22 to 0.50, N = 3; GEB: Mean CGI *r* = 0.50, from 0.32 to 0.61, N = 3). Regarding the comparison networks, although some limited regional overlap existed between the networks of different functions, e.g., SNN and MDN shared 6 regions (Table S1) the GCI scores showed very low correlations (<0.1) across the different functional networks (i.e., between each of the comparison networks with one another, and with any definition of the sentence processing network), which was again consistent across the two independent rs-fMRI datasets for measuring functional connectivity (Fig. 3A; BIL&GIN: Mean *r* = 0.058±0.047, N = 12; GEB: Mean *r* = 0.017±0.12, N = 12).

**Fig. 3.**
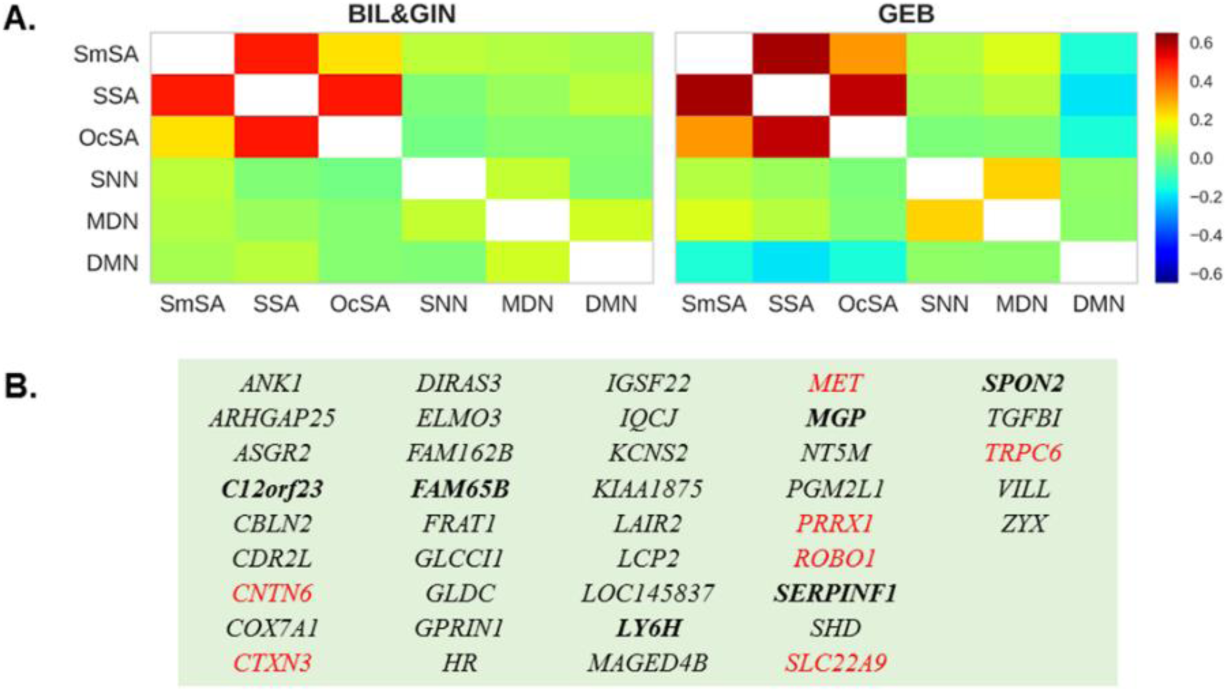
Contributions of individual genes to overall gene-brain network correlations. (A) Correlations between GCI scores derived for each functional network, based on the two independent datasets BIL&GIN (left) and GEB (right). The three definitions of the sentence processing network showed relatively high correlations of their GCI scores, while different functional networks showed low correlations of their GCI scores. (B) The consensus set of genes (*N* = 41) for the six analyses of the sentence processing network (i.e., three functional definition strategies by two rs-fMRI datasets). Red color indicates genes that have been linked to language-related phenotypes in previous studies (e.g., reading deficits, intellectual disability, and autism), and genes **in bold** indicate significant differential expression within the sentence processing network regions compared to other cortical regions (FDR corrected *p* < 0.05). SmSA, Multimodal Sentence Areas; SSA, Synthesized Sentence Areas; OcSA, One-contrast Sentence Areas; SNN, Spatial Navigation Network; MDN, Multiple Demand Network; DMN, Default Mode network.

In order to derive a set of genes related to the sentence processing network with the highest consistency, we identified the 41 “consensus genes” which had positive CGI scores in all six analyses of this network, i.e. the three definition strategies (SmSA, SSA, and OcSA), by the two independent rs-fMRI datasets (BIL&GIN and GEB) (Fig. 3B). As expected, these 41 genes showed significantly greater CGI scores when compared to all other genes, within each of the six analyses (t values > 4.16, *p*s <0.0002; equal variances not assumed). In addition, we re-calculated transcriptomic similarity matrices based on only these 41 consensus genes, and found these matrices to be significantly correlated with the corresponding functional connectivity matrices (*r* = 0.30-0.55, *p*s < 0.0001) (Fig. S4; also the correlations based on the 41 genes were greater than the correlations based on 1000 random gene sets of equal size). These correlations were also largely unaffected after controlling for inter-regional spatial distance between centers of regions (all *p*s < 0.05) (Table S4). Several of the 41 consensus genes, including *ROBO1*, *MET*, *PRRX1*, *CNTN6*, and *CTXN3*, have been reported to affect language- or reading-related phenotypes such as dyslexia, and/or disorders that are often accompanied by linguistic impairments, i.e. intellectual disability, ASD, and schizophrenia (see Discussion). Further information on the consensus gene set is provided in Table S5.

### Biological Roles of Consensus Genes Associated with the Sentence Processing Network

Gene ontology analysis of the consensus set of 41 genes associated with the sentence processing network identified significant enrichment for terms mostly related to the actin cytoskeleton (*p*s < 0.05), all driven by the same set of 5 genes, *MET, ZYX, ARHGAP25, VILL*, and *IGSF22* (Table 1). Another significant enrichment was for the set ‘neuron projection development’ (*p* = 0.012), and there were 6 genes that drove the enrichment results, which were *SERPINF1*, *PRRX1*, *ROBO1*, *TRPC6*, *SPON2*, and *GPRIN1* (Table 1). None of these gene sets was significant when analyzing the 41 top genes for each of the comparison networks (*p* > 0.05) (Table S6).

**Table 1.** Gene ontology terms for which the consensus set associated with the sentence processing network shows enrichment. BP, biological process.

| Term names | Corrected p | # of term genes | Overlap | Intersecting genes |
| --- | --- | --- | --- | --- |
| **BP: neuron projection development** | 0.0118 | 902 | 6 | *SERPINF1*, *PRRX1*, *ROBO1*, *TRPC6*, *SPON2*, *GPRIN1* |
| **BP: actin filament-based process** | 0.0455 | 705 | 5 | *MET*, *ZYX*, *ARHGAP25*, *VILL*, *IGSF22* |
| **BP: actin cytoskeleton organization** | 0.0228 | 609 | 5 | *MET*, *ZYX*, *ARHGAP25*, *VILL*, *IGSF22* |
| **BP: supramolecular fiber organization** | 0.0196 | 590 | 5 | *MET*, *ZYX*, *ARHGAP25*, *VILL*, *IGSF22* |
| **BP: actin filament organization** | 0.00183 | 360 | 5 | *MET*, *ZYX*, *ARHGAP25*, *VILL*, *IGSF22* |

To examine expression across human brain development of these set of genes associated with sentence processing network connectivity, we used data from the Allen Institute’s BrainSpan project, which includes human brain tissues from embryonic stages to adulthood, measured using RNA-sequencing (Li et al., 2018). Each of these genes has detectable expression in the frontal and temporal regions during fetal development and in early childhood (Fig. 4). Some of the genes increase in expression through development all the way to adulthood, such as *MET*, *SERPINF1* and *PRRX1*. Others decrease such as *ROBO1*, *GPRIN1* and *TRPC6*, while still remaining expressed through to adulthood.

**Fig. 4.**
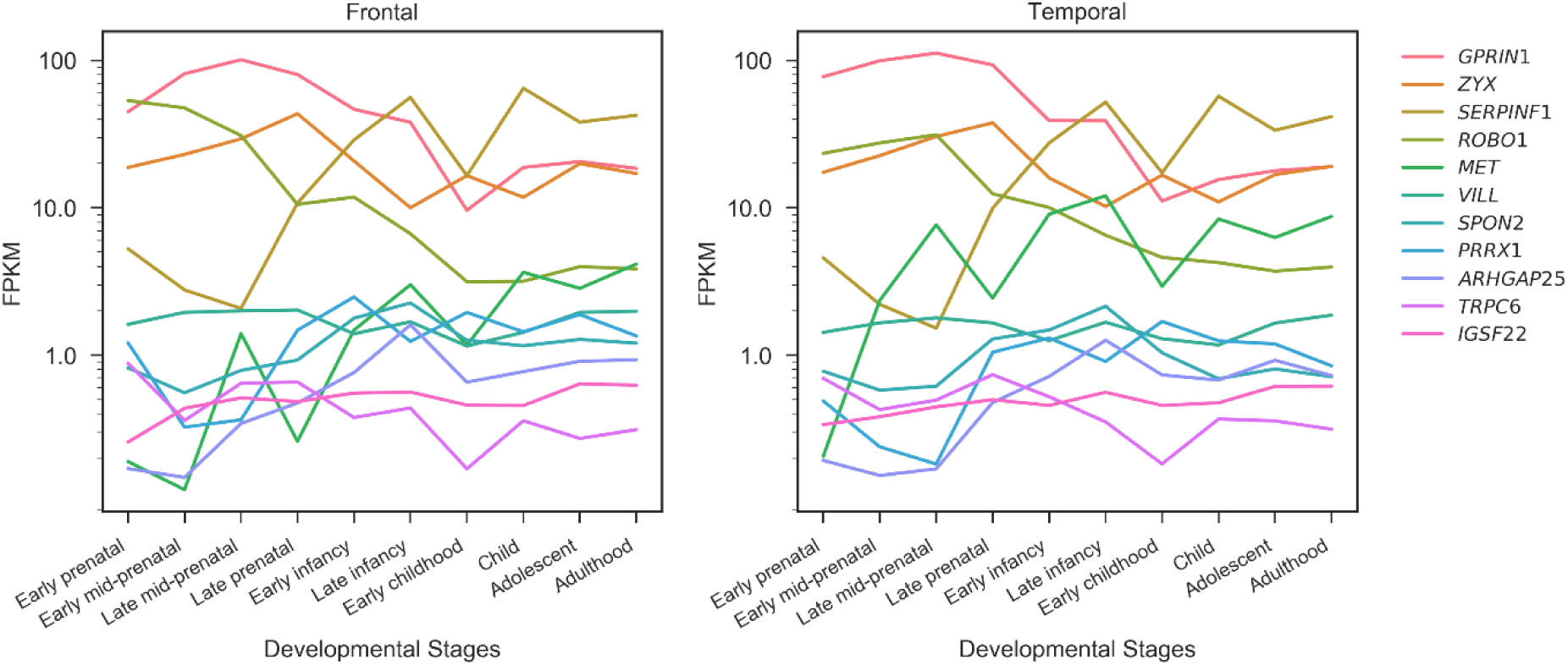
Gene expression in human frontal and temporal cortices, from embryo (8 week post conception) to adulthood (40 years), from the BrainSpan atlas. Genes are listed in the legend in order of their averaged expression across all samples. The y axis uses a log_10_ scale to visualize change over time for each gene. The samples were dissected according to the BrainSpan Technical White Paper (http://www.brainspan.org/).

We found that 14 genes of the 41 consensus gene set showed differential expression when contrasting regions defined as belonging to the sentence processing network against those in the comparison networks (see Methods; uncorrected *p* < 0.05), among which 6 genes survived correction for multiple comparisons (FDR corrected *p* < 0.05): *C12orf23*, *FAM65B*, *LY6H*, *MGP*, *SERPINF1*, and *SPON2*, all with higher expression in the regions assigned to the sentence processing network. Notably, three genes related to “neural projection development” in gene ontology analysis showed significantly higher expression within the sentence-processing regions, which were *SERPINF1*, *SPON2*, and *GPRIN1* (*p*s < 0.05; e.g. *SERPINF1: t*(46) = 3.85, *p* = 0.00036; 95% confidence interval of the difference: 0.24 to 0.76), while none showed lower expression in the sentence processing regions (*p*s >0.05). More information can be found in Table S7.

Of the 41 consensus genes associated with the sentence processing network, data on 33 were available from a single-cell RNA-sequencing study of adult mouse cerebral cortex (Zhang et al., 2014). Among these 33 genes, a majority (i.e., 25) showed expression in one cell type that was at least 1.5 times higher than all other cell types, although not predominantly in neurons versus other cell types (Table S8). For example, *MET* and *CTXN3* showed 3.57- and 1.53-fold expression in neurons compared to the maximum expression in other cell types, respectively. Data on seven of the 41 genes were available in another single-cell gene expression database (Zeisel et al., 2015), and two of these showed enrichment in interneurons: *ANK1* and *SHD*. Among the six genes driving the gene ontology enrichment results related to neural projection development, four showed their highest expressions in neurons, which were *ROBO1*, *TRPC6*, *SPON2*, and *GPRIN1*. In addition, among the five genes driving the gene ontology enrichment related to the actin cytoskeleton, data for three were available: *MET* showed highest expression in neurons, and *ARHGAP25* and *ZYX* in microglia. We also queried several recently available single-cell or single-nucleus gene expression datasets on the human neocortex (Fan et al., 2018; Lake et al., 2018; Li et al., 2018; Zhong et al., 2018). Data on 22 of the 41 genes were available in one or more of these datasets. Among these 22 genes, eight were suggested to be specific to neuronal cell types: *ANK1*, *ROBO1*, *COX7A1*, *LAIR2*, *LY6H*, *GLCCI1*, *CBLN2*, and *PGM2L1* (Table S9). Five of these neuronal genes were markers of excitatory neurons: *ROBO1*, *LAIR2*, *GLCCI1*, *CBLN2*, and *PGM2L1*.

### Association with ASD, schizophrenia and intelligence

We were interested to test whether genes involved in the cortical language network also contain polymorphisms in the population which affect human cognitive or behavioural variation, or susceptibility to neuropsychiatric disorders. This analysis requires genome-wide association scan (GWAS) results from large-scale studies, which are currently lacking for reading/language measures in the general population, and for disorders such as dyslexia or language impairment which involve language-related deficits. We analyzed ASD and schizophrenia, as these disorders can involve linguistic deficits, as well as intelligence in the general population, which correlates with linguistic abilities (see Introduction). Using GWAS summary statistics for ASD based on up to 7387 cases and 8567 controls (Autism Spectrum Disorders Working Group of The Psychiatric Genomics, 2017), we found that the 41 consensus genes associated with the sentence processing network were significantly enriched for single nucleotide polymorphisms (SNPs) showing association with ASD (*beta* = 0.32, *p* = 0.0080, FDR corrected). No such signal in relation to ASD was seen for the top genes (*N* = 41) with highest CGI scores for each of the comparison networks (SNN: *p* = 0.45; MDN: *p* = 0.65; DMN: *p* = 0.074). In addition, no significant enrichment was found for any functional network in relation to schizophrenia (GWAS based on up to 36,989 cases and 113,075 controls; Schizophrenia Working Group of the Psychiatric Genomics, 2014) (*p*s >0.50). For intelligence (GWAS based on 78,308 individuals; (Sniekers et al., 2017)), there was a significant enrichment for the top genes associated with the MDN (*beta* = 0.30, *p* = 0.035), with no other enrichment found (*p*s >0.30).

## Discussion

In this study, we combined gene transcription profiles in the human brain with task and resting-state fMRI data, and investigated the gene expression correlates of the high-level linguistic network. Specifically, with six analyses based on complementary strategies and independent datasets, we revealed a significant correlation between the pattern of functional connectivity within the sentence processing network and the corresponding pattern of inter-regional gene expression similarity. To our knowledge, this is the first evidence for a link between gene transcription profiles and language networks.

While some previous studies have suggested that transcription profiles are linked to patterns of structural and functional connectivity across the brain (see Introduction), this relationship could be driven by broad differences in gene expression between sensory and higher-order association cortices. Here, we focus on one core human cognitive ability – language – and examine a fine-grained pattern of brain-gene relationships within the network that supports sentence processing. Across three definitions of sentence processing regions, and using two independent resting-state fMRI datasets, we established and characterized a relationship with gene expression: pairs of language-sensitive brain regions that show synchronization during rest also show more similar profiles of gene expression. We then identified a consensus set of genes most consistently driving the correlation between transcriptomic similarity and functional connectivity within the sentence processing network, and thereby gained new insights into the molecular bases of the language-ready brain.

The underlying basis of positive correlations between transcriptome similarity and functional connectivity is unknown. One possibility is that functionally linked cortical regions are more likely to share specific aspects of neuronal physiology and developmental trajectory, which support their temporally synchronized activity. When we adjusted for the spatial proximity of regions, the correlations based on 867 genes that comprise the top 5% for differential stability across donors became mostly non-significant, which suggests that these correlations largely reflected local similarities of nearby regions. However, the correlations based on the 41 consensus genes linked to the sentence processing network were not diminished by adjusting for spatial proximity, which indicates that these genes are involved in longer-distance inter-regional relationships within this network.

The consensus set of 41 genes was enriched for functions relating to the actin cytoskeleton, driven by the genes *MET, ZYX, ARHGAP25, VILL*, and *IGSF22*. Mutations in actin cytosleleton-related genes have been tentatively linked to rightward hemispheric language dominance, a rearrangement of language in the brain which occurs in roughly 1% of the population (Carrion-Castillo et al., 2019). The actin cytoskeleton plays many roles in neurons, including in regulating the extension and direction of axon growth (Coles & Bradke, 2015), and the remodeling and maintenance of neuronal architecture throughout the neuron lifetime (Gordon-Weeks & Fournier, 2014). The consensus genes associated with language-related networks were also enriched for roles in neuron projection development. Each of the six genes driving this enrichment, i.e. *SERPINF1*, *PRRX1*, *ROBO1*, *TRPC6*, *SPON2*, and *GPRIN1*, has detectable expression in frontal and temporal regions during fetal development and early childhood, and might play especially important roles during language network development. The fact that these genes, known for their neurodevelopmental roles, are expressed in adult cerebral cortex in a manner linked to functional connectivity within language networks, also suggests that they have continuing roles in maintaining adult cortical circuitry for its regionally-specialized roles. Their continued expression into adulthood certainly attests to adult functions, although these are poorly understood. Further support for the particular importance of these genes for the sentence processing network came from the fact that some of them showed higher gene expression in sentence network regions than elsewhere in the cortex, and none showed significantly lower expression in sentence network regions.

Several of the consensus genes associated with the sentence processing network have previously been reported to contain common polymorphisms or rare mutations which impact on language- and reading-related phenotypes, or else disorders which can be accompanied by reduced linguistic abilities such as intellectual disability, ASD or schizophrenia. *ROBO1* has well established roles in brain development (Hannula-Jouppi et al., 2005; St Pourcain et al., 2014), and has been implicated in dyslexia as well as phonological short term memory (Bates et al., 2011; Hannula-Jouppi et al., 2005), while its homologue *ROBO2* has been linked with expressive vocabulary during early language acquisition (St Pourcain et al., 2014). Mutations in *MET* are a risk factor for ASD (Campbell et al., 2006; Jackson et al., 2009; Mukamel et al., 2011; Sousa et al., 2009), and *MET* is also regulated by the transcription factor *FOXP2* (Mukamel et al., 2011), which causes developmental verbal dyspraxia when mutated (Lai et al., 2001). *PRRX1* is associated with intellectual disability and delayed language acquisition (Lam & Morris, 2016). Mutations of *CNTN6* have been reported in patients with speech and language delays, intellectual disability, and atypical ASD (Kashevarova et al., 2014). *CTXN3* has been linked to schizophrenia (Lewis et al., 2003; Potkin et al., 2010). These observations suggest that our consensus gene set linked to the sentence processing network might provide additional candidates for future studies of language-related individual differences, and neurodevelopmental disorders. In addition, a recent study used functional connectivity based on rs-fMRI, in combination with Allen brain gene expression data, to identify 136 genes highly correlated with the modular organization of intrinsic connectivity networks (although not defined on the basis of task functional data) (Richiardi et al., 2015). We found that nine of those genes overlap with our set of 41 genes (*CNTN6*, *CTXN3*, *LAIR2*, *MGP*, *SHD*, *TGFBI*, *ASGR2*, *CDR2L*, *IQCJ*). The overlap of these genes is encouraging despite the different datasets, methods and brain region definitions used in the other study.

No large-scale GWAS studies have yet been published for reading/language measures in the general population, nor for disorders such as dyslexia or language impairment which involve language-related deficits. However, through analysis of large-scale GWAS summary statistics for ASD, a disorder that can also involve linguistic deficits (Kleinhans et al., 2008; Lombardo et al., 2015), we found that the consensus gene set associated with connectivity in the sentence processing network is enriched for common SNPs that contribute to the polygenic liability to ASD in the population. It is therefore possible that variants in these genes associate with ASD due, at least in part, to dysfunction of high-level language networks. Note that, while language-related deficits may no longer be considered core to ASD in the latest edition of the Diagnostic and Statistical Manual of Mental Disorders (DSM 5) (American Psychiatric Association, 2013), the earlier DSM-IV criteria were used for the large-scale GWAS whose results we used in the present study (Autism Spectrum Disorders Working Group of The Psychiatric Genomics, 2017), in which language delays were considered an important aspect of the disorder (Grzadzinski, Huerta, & Lord, 2013). No such enrichment signal for ASD was observed for gene sets associated with the other functional networks that we analyzed by way of comparison, i.e., spatial navigation, multiple demand, and default mode networks. Interestingly, a recent large-scale brain imaging analysis of ASD found that cortical thinning was present in many of the regions included in our definitions of the sentence-processing network (van Rooij et al., 2018). For schizophrenia and intelligence we found no enrichment of association signals within the consensus gene set associated with the sentence processing network. This pattern may relate to severity, as language deficits in schizophrenia patients have been reported to be less severe than in ASD patients (again based on earlier diagnostic criteria) (Chisholm, Lin, Abu-Akel, & Wood, 2015; Spek & Wouters, 2010). Alternatively, the genetic underpinnings of high-level language processing may be of little relevance to schizophrenia, or intelligence in the general population. However, statistical power to detect these relations may also be an issue.

There is at present no consensus in the field on what should constitute the precise high-level language network and how best to define it (E. Fedorenko & Thompson-Schill, 2014; L. Labache et al., 2018). To circumvent this problem, we focused on sentence-level processing specifically, but used three complementary approaches for defining brain regions important for this function: one based on a conjunction of three task contrasts and functional laterality (SmSA; L. Labache et al., 2018), one based on a large-scale meta-analysis of prior neuroimaging studies (SSA; Yarkoni et al., 2011), and one based on a single task contrast (OcSA; E. Fedorenko et al., 2010). Reassuringly, our three approaches to defining sentence-level processing regions yielded similar sets of regions, especially with respect to areas in the inferior frontal and middle temporal gyri. Likely because of the overlap between the three network definitions, their transcriptomic correlates were also similar, as well as the contributions of individual genes. This overall concordance supports the validity of the different definitions, allowing us to define a consensus set of genes that emerged consistently across all definitions and both of the rs-fMRI datasets, and is thus not dependent solely on any individual approach for defining the functional network.

Although multiple cognitive processes are likely to be involved in sentence processing, at least two of the key elements, syntax and semantics, are tightly integrated in the brain, while combinatorial processing is a key part of both of them (Evelina Fedorenko, Mineroff, Siegelman, & Blank, 2018). Regardless, we do not assume any straightforward mapping of high-level linguistic processing to a small subset of underlying genes. As noted in the introduction, the creation and maintenance of brain circuits underlying language networks surely depends on the coordinated actions of very many genes. Here we show that the overall transcriptional similarity of regions involved in sentence processing is correlated with their functional connectivity, and we identify a relatively small number of genes which contribute to that correlation most strongly and consistently, based on current data. We expect many more genes will be identifiable in relation to this functional network, as higher quality gene expression datasets are generated, and that some combinations of genes may be more or less important for particular cognitive sub-processes.

Functional connectivity of regions during task performance is correlated with that observed at rest, which suggests an “intrinsic” architecture of functional organization (Blank et al., 2014; Cole et al., 2014; L. Labache et al., 2018), which may also be linked to individual differences in behaviour (Arnold, Protzner, Bray, Levy, & Iaria, 2014; Kong, Wang, et al., 2017). Thus, the set of regions for sentence processing defined here could be used in future studies of very large datasets, such as the UK Biobank (Sudlow et al., 2015) or ENIGMA Consortium (Thompson et al., 2014), which include neuroanatomical and/or intrinsic connectivity data but limited task fMRI data for language functions. For instance, the functional regions defined here could be used to investigate structural/functional variability during maturation, aging, and/or pathological processes in the general population, including those associated with developmental language disorders.

For the comparison networks included in this study, i.e. the spatial navigation, multiple demand, and default mode networks, we also found similar overall correlations between the functional connectivity networks and the corresponding transcriptomic networks, although with largely different individual genes contributing. This was indicated by relatively low correlations of GCI scores across different networks, and also that different functional networks produced distinct results in gene ontology analysis. There were individual genes such as *MET* that showed a relatively high contribution to the connectivity-transcriptomic correlation for the sentence processing network, but negligible contributions to other functional networks. Therefore, although the existence of connectivity-transcriptome correlations appears to be ubiquitous across functional networks, our study yields the important basic insight that different functional networks can involve differently weighted genetic contributions at the level of cortical gene expression. This is broadly consistent with genetic correlations between different cognitive abilities such as verbal and non-verbal performance, as assessed in population genetic analysis, which often indicate shared but also independent genetic effects on such pairs of traits (Trzaskowski, Shakeshaft, & Plomin, 2013). Our study therefore suggests a novel approach to complement existing genetic epidemiological approaches, for understanding the general versus specific influences on diverse cognitive abilities. Note that, in the present study, in order to achieve comparable statistical power in the network similarity analysis across different networks, we purposely defined similar numbers of top regions for each comparison network, even though this meant using different thresholds for including regions in each comparison network. Further studies may investigate how using more or less inclusive definitions of these functional networks affect relationships with gene expression.

The present study was based on gene expression data from a small number of donors in the Allen Brain database. Post-mortem brain tissues suitable for transcriptomic analysis are difficult to collect from individuals who were healthy immediately prior to death, which is necessary since RNA degrades within hours after death. Thus, the availability of high-quality gene expression data from the human brain is necessarily limited. Moreover, most of the Allen brain data, and all the data used for the present study, are based on the older and relatively noisy transcriptomic technology of microarrays, rather than the more accurate, latest method of RNA sequencing. In addition, some genes known to be involved in language, such as *FOXP2* (Lai et al., 2001), had to be excluded from our analyses because of the inclusion criterion of stability across donors. *FOXP2* showed a low inter-donor differential stability of 0.22 across cerebral cortex samples, especially low across frontal samples (−0.07) and across temporal samples (0.16) in the Allen brain data (M. Hawrylycz et al., 2015). Thus, it is likely that data of sufficient quality were not available for other genes too, that might have been of relevance to language networks, such that future studies using RNA sequencing in larger numbers of individuals, and with more sampling per cortical region, would be well motivated. In addition, inter-individual variabilities have been observed in the precise locations of language-sensitive regions (E. Fedorenko et al., 2010; Steinmetz & Seitz, 1991) as well as the cytoarchitectonic features of higher level cognitive areas (e.g., BA 44 and 45) (Fischl et al., 2008). To address these issues, we focused on only a subset of genes with the highest differential stability across donors, in order to ensure a reliable and representative, average map of gene expression, which was necessary for this study. Inter-donor differential stability is particularly low for cortical regions in the Allen dataset, for the majority of individual genes (Fig. S2), so that only a minority of genes have a useful degree of consistency for group-based analysis. Having made a selection based on high stability across donors, the reproducibility of our genetic findings with respect to two independent rs-fMRI datasets supported their validity. Gene expression data from a larger number of individuals, ideally when both anatomical and functional data from the same individuals are available, would help to further improve upon this aspect. However, the prospective enrollment of eventual donors of post mortem tissue into brain imaging studies during life would be extremely difficult to achieve, within a typical project duration.

As noted in the Introduction, several recent studies have examined the relationship between post mortem brain transcriptomics and in vivo neuroimaging in different datasets (e.g., Anderson et al., 2018; Reardon et al., 2018; Richiardi et al., 2015; Seidlitz et al., 2018; Whitaker et al., 2016). While these studies have provided strong evidence that patterns of gene expression co-vary with anatomical and functional organization of the human brain, none of them focused on specific core cognitive functions such as language. Unlike these previous studies, the present study provided new evidence for the gene expression correlates of a high-level language network, and additionally identified a set of individual genes likely to be important for cortical language functions.

A limitation of our study is the use of transcriptomic data from blocks of cerebral cortical tissues, which comprised many cell types. Future databases based on single-cell transcriptomics would likely provide further insights. Although efforts to produce such data are underway for a limited number of human cerebral cortical regions, no database currently exists which has broad mapping over the cerebral cortex. It will be a major undertaking in the future for the human brain science community to achieve a widespread cerebral cortical gene expression map at single cell resolution. For the time being, we queried our consensus genes associated with the language-related network using single-cell or single-nucleus transcriptomic data from brain tissues from a restricted number of regions/structures, which gave information about whether specific genes are relatively more highly expressed in neurons versus some major classes of glial cells. However, even if a gene of interest was expressed at a comparable or higher level in glia than neurons, it might still influence neuronal physiology and circuit properties, either directly through its expression in neurons, or indirectly via the interactions of surrounding glial cells with neurons.

As regards head motion during scanning, both signal artifacts (e.g., Friston, Williams, Howard, Frackowiak, & Turner, 1996; Kong, 2014; Power, Barnes, Snyder, Schlaggar, & Petersen, 2012; Satterthwaite et al., 2012) and functionally meaningful and reliable variability (e.g., Couvy-Duchesne et al., 2014; Hodgson et al., 2017; Kong et al., 2014; Zeng et al., 2014) have been suggested to relate to this. In the present study, we carried out traditional preprocessing in relation to head motion, including motion correction, quality control in terms of the amount of head motion, and confound regression, but we did not apply further preprocessing steps proposed more recently, such as the “scrubbing” method (Power et al., 2012). Motion issues mainly affect data from young children, the elderly, and disorder groups (Kong et al., 2014; Satterthwaite et al., 2012), while most of the subjects involved in the present study were healthy young adults, with a good capacity to stay still in scanners. More importantly, we found high correlations (up to 0.92) of the functional connectivity patterns extracted from two independent rs-fMRI datasets (i.e., BIL&GIN and GEB), and we focused on consensus genes arising from the analysis of these two datasets. Thus, any motion-induced artifacts would have minimal impact on our analyses.

In sum, we provide a first description of the overall transcriptomic correlates of brain networks underlying high-level linguistic processing, as well as identifying a set of individual genes likely to be most important. These findings help elucidate the molecular basis of language networks, as distinct from functional networks important for other aspects of cognition. A link of this genetic infrastructure to ASD is also suggested by our data. Finally, we propose functional connectivity and gene expression analysis as a complementary approach to existing genetic epidemiological and genetic association approaches, for understanding complex cognitive traits.

## Acknowledgements

We thank the NeuroSynth, Allen Human Brain Atlas, and the Psychiatric Genomics Consortium for data sharing. This study was funded by the Max Planck Society (Germany).

