## Supplementary material for "Gene Expression Correlates of the Cortical Network Underlying Sentence Processing": Table S

**Table S1. Left hemisphere AICHA atlas regions included in each functional network.** See main text for the meanings of the network abbreviations. Region labels in *italic* indicate regions assigned to the language-related network according to either definition, i.e. SmSA, SSA and OcSA.

| ID | Label | SmSA | SSA | OcSA | SNN | MDN | DMN |
|---|---|---|---|---|---|---|---|
| *3* | *G_Frontal_Sup-2-L* | YES | | | | | YES |
| *29* | *S_Inf_Frontal-1-L* | | YES | | | | |
| *31* | *S_Inf_Frontal-2-L* | YES | YES | | | YES | |
| *33* | *G_Frontal_Inf_Tri-1-L* | YES | YES | YES | | | |
| *41* | *G_Frontal_Inf_Orb-1-L* | YES | | | | | |
| *57* | *S_Precentral-4-L* | YES | YES | YES | | | |
| *99* | *G_SupraMarginal-7-L* | YES | YES | YES | | | |
| *103* | *G_Angular-2-L* | YES | | | | | YES |
| *105* | *G_Angular-3-L* | | | YES | | | YES |
| *111* | *S_Intraparietal-2-L* | | YES | | | YES | |
| *141* | *G_Occipital_Inf-1-L* | YES | | | | | |
| *147* | *G_Insula-anterior-2-L* | YES | | | | | |
| *149* | *G_Insula-anterior-3-L* | YES | YES | | | | |
| *151* | *G_Insula-anterior-4-L* | | YES | | | | |
| *165* | *G_Temporal_Sup-3-L* | | YES | | | | |
| *167* | *G_Temporal_Sup-4-L* | YES | YES | YES | | | |
| *169* | *S_Sup_Temporal-1-L* | YES | YES | YES | | | |
| *171* | *S_Sup_Temporal-2-L* | YES | YES | YES | | | |
| *173* | *S_Sup_Temporal-3-L* | YES | YES | YES | | | |
| *175* | *S_Sup_Temporal-4-L* | YES | YES | YES | | | |
| *177* | *S_Sup_Temporal-5-L* | | YES | YES | | | YES |
| *179* | *G_Temporal_Mid-1-L* | | YES | YES | | | YES |
| *183* | *G_Temporal_Mid-3-L* | YES | YES | YES | | | |
| *185* | *G_Temporal_Mid-4-L* | YES | YES | | | | |
| *193* | *G_Temporal_Inf-4-L* | YES | YES | | | YES | |
| *223* | *G_Supp_Motor_Area-2-L* | YES | | | | | |
| *225* | *G_Supp_Motor_Area-3-L* | YES | YES | | | | |
| *229* | *S_Cingulate-2-L* | | YES | | | | |
| *263* | *G_Paracentral_Lobule-4-L* | YES | | | | | |
| *275* | *G_Precuneus-6-L* | YES | | | YES | YES | |
| *337* | *G_Fusiform-4-L* | | YES | | | | |
| *13* | S_Sup_Frontal-4-L | | | | | | YES |
| *17* | S_Sup_Frontal-6-L | | | | YES | YES | |
| *27* | G_Frontal_Mid-5-L | | | | | YES | |
| *53* | S_Precentral-2-L | | | | | YES | |
| *81* | G_Parietal_Sup-3-L | | | | | YES | |

| | | | | |
|---|---|---|---|---|
| *83* | G_Parietal_Sup-4-L | | YES | |
| *85* | G_Parietal_Sup-5-L | YES | YES | |
| *109* | S_Intraparietal-1-L | | YES | |
| *113* | S_Intraparietal-3-L | | YES | |
| *127* | G_Occipital_Lat-5-L | | YES | |
| *131* | G_Occipital_Sup-2-L | | | |
| *133* | G_Occipital_Mid-1-L | YES | YES | |
| *135* | G_Occipital_Mid-2-L | | YES | |
| *139* | G_Occipital_Mid-4-L | YES | | |
| *207* | G_Frontal_Sup_Medial-1-L | | | YES |
| *209* | G_Frontal_Sup_Medial-2-L | | | YES |
| *213* | S_Anterior_Rostral-1-L | | | YES |
| *217* | G_Frontal_Med_Orb-2-L | | | YES |
| *265* | G_Precuneus-1-L | YES | | |
| *267* | G_Precuneus-2-L | | | YES |
| *269* | G_Precuneus-3-L | | | YES |
| *273* | G_Precuneus-5-L | YES | | |
| *279* | G_Precuneus-8-L | YES | | |
| *283* | S_Parietooccipital-1-L | YES | | |
| *287* | S_Parietooccipital-3-L | YES | | |
| *289* | S_Parietooccipital-4-L | YES | | |
| *293* | S_Parietooccipital-6-L | YES | | |
| *299* | G_Calcarine-1-L | YES | | |
| *305* | G_Lingual-1-L | YES | | |
| *309* | G_Lingual-3-L | YES | | |
| *323* | G_ParaHippocampal-2-L | YES | | |
| *329* | G_ParaHippocampal-5-L | YES | | |
| *341* | G_Fusiform-6-L | YES | YES | |
| *343* | G_Fusiform-7-L | YES | YES | |

**Table S2. Overall correlations between the functional networks and their corresponding transcriptomic networks, based on the two resting state datasets BIL&GIN and GEB, and different sets of genes defined by their differential stability across donors in the Allen brain data (see Methods).** Spearman correlations in bold indicate being significant ($p < 0.05$).

| | | **SMSA** | **SSA** | **OCSA** | **SNN** | **MDN** | **DMN** |
|---|---|---|---|---|---|---|---|
| **TOP 5%** | BIL&GIN | **0.19** | **0.26** | **0.42** | **0.24** | **0.39** | **0.4** |
| | GEB | **0.19** | **0.24** | **0.35** | **0.33** | **0.5** | **0.6** |
| **TOP 10%** | BIL&GIN | **0.16** | **0.23** | **0.36** | **0.22** | **0.37** | **0.3** |
| | GEB | **0.15** | **0.21** | **0.26** | **0.31** | **0.46** | **0.54** |
| **BOTTOM 5%** | BIL&GIN | 0.05 | -0.002 | 0.14 | **0.23** | 0.05 | -0.01 |
| | GEB | 0.02 | -0.005 | 0.02 | 0.14 | 0.05 | 0.15 |
| **BOTTOM 10%** | BIL&GIN | 0.06 | -0.0006 | 0.16 | **0.23** | 0.04 | 0.0004 |
| | GEB | 0.03 | -0.003 | 0.05 | 0.15 | 0.03 | 0.16 |

**Table S3. Overall correlations between the functional networks and their corresponding transcriptomic networks, after controlling for inter-regional spatial distance, based on the top 5% of genes by differential stability across donors in the Allen brain data (see Methods).**

| | **BIL&GIN** | | **GEB** | |
|---|---|---|---|---|
| | **rho** | ***p*** | **rho** | ***p*** |
| **SMSA** | 0.18 | 0.024 | 0.06 | 0.373 |
| **SSA** | 0.16 | 0.036 | 0.07 | 0.307 |
| **OCSA** | 0.25 | 0.043 | 0.06 | 0.628 |
| **SNN** | 0.14 | 0.089 | 0.13 | 0.090 |
| **MDN** | 0.11 | 0.258 | 0.01 | 0.935 |
| **DMN** | 0.02 | 0.847 | 0.19 | 0.125 |

**Table S4. Correlations between the functional networks and their corresponding transcriptomic networks, after controlling for inter-regional spatial distance, based only on the 41 consensus genes linked to the sentence processing network (see Methods).** Spearman correlations in bold indicate being significant ($p < 0.05$).

| | **BIL&GIN** | | **GEB** | |
|---|---|---|---|---|
| | **rho** | ***p*** | **rho** | ***p*** |
| **SMSA** | **0.14** | 0.048 | **0.17** | 0.017 |
| **SSA** | **0.21** | 0.0023 | **0.14** | 0.038 |
| **OCSA** | **0.41** | 0.0011 | **0.40** | 0.0018 |
| **SNN** | -0.02 | 0.77 | -0.02 | 0.81 |
| **MDN** | -0.13 | 0.15 | -0.23 | 0.0099 |
| **DMN** | 0.18 | 0.17 | -0.21 | 0.11 |

**Table S5. The consensus gene set linked to connectivity within the sentence processing network.** GCI1-6 indicates the gene contribution index scores in the six analyses for the sentence processing network.

| GENE | GENE_NAME | ENTREZ_ID | CHR. | GCI1 ($10^{-3}$) | GCI2 ($10^{-3}$) | GCI3 ($10^{-3}$) | GCI4 ($10^{-3}$) | GCI5 ($10^{-3}$) | GCI6 ($10^{-3}$) |
|---|---|---|---|---|---|---|---|---|---|
| **ANK1** | ankyrin 1, erythrocytic | 286 | 8 | 0.27 | 0.51 | 0.23 | 0.61 | 0.22 | 0.27 |
| **ASGR2** | asialoglycoprotein receptor 2 | 433 | 17 | 0.54 | 0.63 | 0.94 | 2.32 | 0.58 | 0.82 |
| **COX7A1** | cytochrome c oxidase subunit VIIa polypeptide 1 (muscle) | 1346 | 19 | 0.68 | 0.72 | 0.19 | 0.94 | 0.03 | 0.07 |
| **GLDC** | glycine dehydrogenase (decarboxylating) | 2731 | 9 | 0.16 | 0.23 | 0.36 | 0.07 | 0.12 | 0.20 |
| **KCNS2** | potassium voltage-gated channel, delayed-rectifier, subfamily S, member 2 | 3788 | 8 | 0.21 | 0.19 | 0.36 | 0.07 | 0.01 | 0.08 |
| **LAIR2** | leukocyte-associated immunoglobulin-like receptor 2 | 3904 | 19 | 1.28 | 2.48 | 2.40 | 4.03 | 0.86 | 2.35 |
| **LCP2** | lymphocyte cytosolic protein 2 (SH2 domain containing leukocyte protein of 76kDa) | 3937 | 5 | 0.07 | 0.17 | 0.07 | 1.99 | 0.18 | 0.61 |
| **LY6H** | lymphocyte antigen 6 complex, locus H | 4062 | 8 | 0.23 | 0.18 | 0.36 | 0.07 | 0.23 | 0.05 |
| **MET** | met proto-oncogene (hepatocyte growth factor receptor) | 4233 | 7 | 1.27 | 3.79 | 0.53 | 5.28 | 3.77 | 6.14 |
| **MGP** | matrix Gla protein | 4256 | 12 | 2.40 | 3.08 | 0.48 | 1.69 | 3.37 | 4.39 |
| **SERPINF1** | serpin peptidase inhibitor, clade F (alpha-2 antiplasmin, pigment epithelium derived factor), member 1 | 5176 | 17 | 0.04 | 0.10 | 0.36 | 0.07 | 0.18 | 0.14 |
| **PRRX1** | paired related homeobox 1 | 5396 | 1 | 0.41 | 0.47 | 1.07 | 1.78 | 0.18 | 0.35 |
| **ROBO1** | roundabout, axon guidance receptor, homolog 1 (Drosophila) | 6091 | 3 | 0.19 | 0.33 | 0.36 | 0.07 | 0.05 | 0.06 |
| **TGFBI** | transforming growth factor, beta-induced, 68kDa | 7045 | 5 | 1.43 | 0.60 | 1.99 | 3.57 | 2.54 | 1.37 |
| **TRPC6** | transient receptor potential cation channel, subfamily C, member 6 | 7225 | 11 | 0.16 | 0.09 | 0.19 | 0.94 | 0.03 | 0.07 |
| **ZYX** | zyxin | 7791 | 7 | 0.07 | 0.22 | 0.23 | 0.61 | 0.19 | 0.25 |
| **DIRAS3** | DIRAS family, GTP-binding RAS-like 3 | 9077 | 1 | 0.10 | 0.15 | 1.78 | 1.90 | 0.80 | 0.80 |
| **FAM65B** | family with sequence similarity 65, member B | 9750 | 6 | 0.02 | 0.08 | 0.36 | 0.07 | 0.39 | 0.23 |
| **ARHGAP25** | Rho GTPase activating protein 25 | 9938 | 2 | 0.16 | 0.12 | 0.32 | 0.40 | 0.55 | 0.56 |
| **FRAT1** | frequently rearranged in advanced T-cell lymphomas | 10023 | 10 | 0.11 | 0.31 | 0.19 | 0.94 | 0.16 | 0.20 |
| **SPON2** | spondin 2, extracellular matrix protein | 10417 | 4 | 1.04 | 1.12 | 0.23 | 0.61 | 0.04 | 0.10 |
| **CNTN6** | contactin 6 | 27255 | 3 | 0.24 | 0.76 | 0.32 | 1.74 | 0.30 | 0.84 |
| **CDR2L** | cerebellar degeneration-related protein 2-like | 30850 | 17 | 0.18 | 0.11 | 0.36 | 0.07 | 0.16 | 0.11 |
| **VILL** | villin-like | 50853 | 3 | 0.06 | 0.08 | 0.23 | 0.61 | 0.29 | 0.21 |
| **HR** | hairless homolog (mouse) | 55806 | 8 | 0.20 | 0.18 | 0.36 | 0.07 | 0.16 | 0.20 |

| | | | | | | | | | |
|---|---|---|---|---|---|---|---|---|---|
| **NT5M** | 5',3'-nucleotidase, mitochondrial | 56953 | 17 | 0.07 | 0.09 | 0.36 | 0.07 | 0.06 | 0.10 |
| **SHD** | Src homology 2 domain containing transforming protein D | 56961 | 19 | 0.02 | 0.36 | 0.36 | 0.07 | 0.61 | 0.74 |
| **ELMO3** | engulfment and cell motility 3 | 79767 | 16 | 1.15 | 0.82 | 0.94 | 2.32 | 0.21 | 0.34 |
| **MAGED4B** | melanoma antigen family D, 4B | 81557 | X | 0.17 | 0.21 | 1.07 | 1.78 | 0.03 | 0.21 |
| **C12ORF23** | chromosome 12 open reading frame 23 | 90488 | 12 | 0.18 | 0.16 | 0.36 | 0.07 | 0.04 | 0.05 |
| **GLCCI1** | glucocorticoid induced transcript 1 | 113263 | 7 | 0.13 | 0.07 | 0.36 | 0.07 | 0.15 | 0.08 |
| **SLC22A9** | solute carrier family 22 (organic anion transporter), member 9 | 114571 | 11 | 0.41 | 0.59 | 0.19 | 1.44 | 0.75 | 1.24 |
| **GPRIN1** | G protein regulated inducer of neurite outgrowth 1 | 114787 | 5 | 0.23 | 0.18 | 0.36 | 0.07 | 0.08 | 0.02 |
| **LOC145837** | uncharacterized LOC145837 | 145837 | 15 | 0.11 | 0.14 | 0.36 | 0.07 | 0.04 | 0.06 |
| **CBLN2** | cerebellin 2 precursor | 147381 | 18 | 0.26 | 0.55 | 1.07 | 1.78 | 0.82 | 1.19 |
| **FAM162B** | family with sequence similarity 162, member B | 221303 | 6 | 0.11 | 0.15 | 0.36 | 0.07 | 0.01 | 0.10 |
| **PGM2L1** | phosphoglucomutase 2-like 1 | 283209 | 11 | 0.06 | 0.14 | 0.36 | 0.07 | 0.17 | 0.08 |
| **IGSF22** | immunoglobulin superfamily, member 22 | 283284 | 11 | 0.19 | 0.17 | 0.36 | 0.07 | 0.06 | 0.08 |
| **KIAA1875** | KIAA1875 | 340390 | 8 | 0.26 | 0.74 | 1.07 | 1.78 | 0.35 | 0.70 |
| **CTXN3** | cortexin 3 | 613212 | 5 | 3.08 | 2.09 | 5.33 | 6.70 | 5.85 | 5.54 |
| **IQCJ** | IQ motif containing J | 654502 | 3 | 0.37 | 0.34 | 0.36 | 0.07 | 0.29 | 0.22 |

**Table S6. Gene ontology terms for the top genes driving connectivity-transcriptomic correlations in the comparison networks.**

| NETWORK | GENE ONTOLOGY TERM | CORRECTED P VALUE | TERMS | OVERLAP | INTERSECTING GENES |
|---|---|---|---|---|---|
| **SNN** | BP: regulation of cellular response to growth factor stimulus | 0.0111 | 259 | 4 | ADAMTS3, ONECUT2, ONECUT1, IL1B |
| | BP: regulation of cell adhesion | 0.0225 | 628 | 5 | FES, SKAP1, ONECUT2, ONECUT1, IL1B |
| | BP: taxis | 0.0157 | 582 | 5 | LHX9, SPON2, FES, FEZF1, IL1B |
| **MDN** | BP: response to antibiotic | 0.0382 | 335 | 4 | PENK, MET, OPRM1, RBP4 |
| | BP: axon development | 0.00822 | 478 | 5 | TNN, SPON2, FEZF1, LHX9, EPHA3 |
| | BP: cellular response to oxidative stress | 0.0199 | 283 | 4 | PENK, MET, MGST1, GPX3 |
| | BP: neuropeptide signaling pathway | 0.023 | 101 | 3 | PENK, OPRM1, PDYN |
| | BP: regulation of production of molecular mediator of immue response | 0.0338 | 115 | 3 | SPON2, KLK7, EBP4 |
| | BP: positive regulation of production of molecular mediator of immune response | 0.00941 | 75 | 3 | SPON2, KLK7, EBP4 |
| | BP: response to lipid | 0.0142 | 907 | 6 | PENK, SPON2, OPRM1, EPHA3, RBP4, MGST1 |
| | BP: response to bacterium | 0.0221 | 588 | 5 | PENK, SPON2, OPRM1, KLK7, MGST1 |
| | BP: cell morphogenesis involved in differentiation | 0.0432 | 678 | 5 | TNN, MET, SPON2, FEZF1, LHX9 |
| | BP: synaptic signaling | 0.0255 | 606 | 5 | PENK, OPRM1, BAIAP3, PDYN, CHRNA2 |
| | BP: response to drug | 0.0196 | 961 | 6 | PENK, MET, OPRM1, RBP4, MGST1, CHRNA2 |
| | BP: response to toxic substance | 0.0000181 | 498 | 7 | PENK, MET, OPRM1, RBP4, MGST1, GXP1, CHRNA2 |
| | CC: synapse | 0.000638 | 846 | 7 | PENK, OPRM1, SV2C, BAIAP3, COL13A1, PDYN, CHRNA2 |
| **DMN** | BP: axon development | 0.011 | 478 | 4 | SCN1B, NEFH, TSPAN2, NEFM, TNN |
| | BP: regulation of Wnt signaling pathway | 0.0447 | 328 | 4 | SIX3, SOST, TNN, ANKRD6 |
| | BP: negative regulation of Wnt signaling pathway | 0.00677 | 202 | 4 | SIX3, SOST, TNN, ANKRD6 |
| | BP: regulation of ossification | 0.00497 | 185 | 4 | SOST, TWIST2, MGP, TNN |
| | CC: transporter complex | 0.0426 | 324 | 4 | SCN1B, GABRA5, GABRQ, KCNS1 |
| | CC: extracellular matrix | 0.0234 | 560 | 5 | SOST, COCH, FREM3, MGP, TNN |
| | CC: axon | 0.0166 | 521 | 5 | SCN1B, PDYN, NEFH, NECAB2, NEFM |
| | MF: gated channel activity | 0.0491 | 336 | 4 | SCN1B, GABRA5, GABRQ, KCNS1 |
| | MF: ion gated channel activity | 0.0474 | 333 | 4 | SCN1B, GABRA5, GABRQ, KCNS1 |
| | MF: structural constituent of cytoskeleton | 0.0339 | 108 | 3 | KRT31, NEFH, NEFM |

BP, biological process; CC: cellular component; MF: molecular function. SNN, Spatial Navigation Network; MDN, Multiple Demands Network; DMN, Default Mode Network

**Table S7. Preferential expression of the consensus set of genes linked to sentence processing network connectivity, when contrasting sentence processing regions against regions of the comparison networks.**

| GENE | T | DF | SIG. (2-TAILED) | MEAN DIFFERENCE | STD. ERROR DIFFERENCE | 95% CI OF THE DIFFERENCE | | FDR_P |
|---|---|---|---|---|---|---|---|---|
| ANK1 | -2.275 | 61.585 | 0.026378 | -0.29684 | 0.130462 | -0.55767 | -0.03602 | 0.098319 |
| ASGR2 | 1.264 | 59.608 | 0.211 | 0.1527 | 0.120766 | -0.0889 | 0.394301 | 0.286985 |
| COX7A1 | 0.262 | 59.983 | 0.793969 | 0.033411 | 0.127364 | -0.22136 | 0.288178 | 0.793969 |
| GLDC | 1.716 | 53.484 | 0.091937 | 0.151093 | 0.088046 | -0.02547 | 0.327654 | 0.169536 |
| KCNS2 | 1.373 | 40.608 | 0.177323 | 0.138121 | 0.100606 | -0.06512 | 0.341358 | 0.269268 |
| LAIR2 | 1.982 | 56.66 | 0.052383 | 0.348645 | 0.17594 | -0.00372 | 0.701005 | 0.124933 |
| LCP2 | 1.827 | 62.881 | 0.072423 | 0.226678 | 0.124061 | -0.02125 | 0.474602 | 0.156282 |
| LY6H | 4.036 | 52.416 | 0.000178 | 0.518619 | 0.12849 | 0.260834 | 0.776404 | 0.003819 |
| MET | -0.592 | 60.243 | 0.556127 | -0.15925 | 0.26905 | -0.69739 | 0.378882 | 0.584646 |
| MGP | 2.874 | 53.879 | 0.005797 | 0.545522 | 0.189843 | 0.16489 | 0.926155 | 0.039611 |
| SERPINF1 | 3.851 | 45.857 | 0.000363 | 0.499352 | 0.129672 | 0.238313 | 0.760391 | 0.003819 |
| PRRX1 | -1.701 | 57.254 | 0.094331 | -0.13388 | 0.0787 | -0.29146 | 0.023694 | 0.169536 |
| ROBO1 | 1.695 | 61.486 | 0.095105 | 0.108899 | 0.064241 | -0.01954 | 0.237337 | 0.169536 |
| TGFBI | 1.248 | 59.299 | 0.216989 | 0.241808 | 0.193779 | -0.1459 | 0.629518 | 0.286985 |
| TRPC6 | 1.47 | 61.16 | 0.146672 | 0.124961 | 0.085004 | -0.04501 | 0.294928 | 0.240542 |
| ZYX | -0.845 | 63.94 | 0.401502 | -0.0827 | 0.097918 | -0.27831 | 0.112919 | 0.448392 |
| DIRAS3 | 2.646 | 62.554 | 0.010294 | 0.206882 | 0.078192 | 0.050607 | 0.363158 | 0.052756 |
| FAM65B | 3.876 | 56.99 | 0.000277 | 0.475978 | 0.122814 | 0.230046 | 0.721911 | 0.003819 |
| ARHGAP25 | 1.956 | 63.696 | 0.054849 | 0.220125 | 0.112536 | -0.00471 | 0.444961 | 0.124933 |
| FRAT1 | -2.162 | 61.386 | 0.034562 | -0.20808 | 0.096265 | -0.40055 | -0.01561 | 0.099955 |
| SPON2 | 3.782 | 57.48 | 0.000373 | 0.619676 | 0.163847 | 0.291638 | 0.947714 | 0.003819 |
| CNTN6 | 1.414 | 58.89 | 0.162508 | 0.161233 | 0.113993 | -0.06688 | 0.389343 | 0.256262 |
| CDR2L | -2.026 | 60.661 | 0.047118 | -0.18518 | 0.091381 | -0.36793 | -0.00243 | 0.12074 |
| VILL | -1.143 | 63.961 | 0.257482 | -0.0705 | 0.061702 | -0.19376 | 0.052767 | 0.329898 |
| HR | -1.802 | 62.44 | 0.076358 | -0.14571 | 0.080854 | -0.30731 | 0.015895 | 0.156533 |
| NT5M | -1.289 | 62.833 | 0.202002 | -0.09297 | 0.072107 | -0.23707 | 0.05113 | 0.28559 |
| SHD | -0.938 | 62.914 | 0.351778 | -0.1257 | 0.133994 | -0.39347 | 0.142073 | 0.424203 |
| ELMO3 | -1.299 | 63.999 | 0.198597 | -0.16419 | 0.126393 | -0.41669 | 0.088311 | 0.28559 |
| MAGED4B | 2.666 | 62.135 | 0.009776 | 0.183261 | 0.068745 | 0.045847 | 0.320674 | 0.052756 |
| C12ORF23 | 3.269 | 64 | 0.001738 | 0.197655 | 0.06046 | 0.076873 | 0.318437 | 0.01425 |
| GLCCI1 | -1.063 | 63.122 | 0.291919 | -0.09197 | 0.086539 | -0.2649 | 0.080954 | 0.362688 |
| SLC22A9 | 1.672 | 62.007 | 0.099652 | 0.221549 | 0.132541 | -0.0434 | 0.486493 | 0.170239 |
| GPRIN1 | 2.194 | 63.8 | 0.031874 | 0.109142 | 0.049742 | 0.009764 | 0.208519 | 0.099955 |
| LOC145837 | 0.649 | 56.511 | 0.518946 | 0.03438 | 0.05297 | -0.07171 | 0.140471 | 0.559915 |
| CBLN2 | 2.468 | 50.079 | 0.017038 | 0.395125 | 0.160086 | 0.073595 | 0.716654 | 0.077616 |
| FAM162B | 0.84 | 56.843 | 0.404647 | 0.09039 | 0.107658 | -0.1252 | 0.305985 | 0.448392 |
| PGM2L1 | 2.362 | 63.853 | 0.021263 | 0.126594 | 0.053608 | 0.019496 | 0.233692 | 0.08718 |
| IGSF22 | 2.204 | 63.996 | 0.031107 | 0.154815 | 0.070233 | 0.014507 | 0.295122 | 0.099955 |
| KIAA1875 | -2.136 | 63.417 | 0.036569 | -0.20559 | 0.096263 | -0.39793 | -0.01325 | 0.099955 |
| CTXN3 | 0.517 | 63.921 | 0.607064 | 0.130138 | 0.251805 | -0.37291 | 0.633188 | 0.622241 |
| IQCJ | 0.886 | 63.957 | 0.378901 | 0.075186 | 0.084854 | -0.09433 | 0.244703 | 0.443856 |

**Table S8. Preferential expression in specific cell-types of the consensus set of genes linked to sentence processing network connectivity.** The cell-type specificity data were from mouse cortex (Zhang et al., 2014; Zeisel et al., 2015).

| GENE | CHR. | ASTROCYTES | NEURON | OLIGODENDROCYTE PRECURSOR CELL | NEWLY FORMED OLIGODENDROCYTE | MYELINATING OLIGODENDROCYTES | MICROGLIA | ENDOTHELIAL CELLS | ZHANG_ENRICH | ZHANG_ENRICH2 | ZHANG2014 | ZEISEL2015 |
|---|---|---|---|---|---|---|---|---|---|---|---|---|
| **ANK1** | 8 | 0.228 | 4.395 | 3.691 | 1.227 | 0.340 | 0.214 | 0.100 | 1.191 | 3.583 | | Interneuron |
| **ASGR2** | 17 | 0.100 | 0.100 | 0.100 | 0.298 | 0.129 | 1.318 | 0.100 | 4.429 | 10.255 | Microglia | |
| **COX7A1** | 19 | 1.514 | 3.257 | 0.688 | 0.379 | 1.215 | 0.397 | 0.686 | 2.151 | 2.682 | Neuron | Ependymal |
| **GLDC** | 9 | 25.924 | 1.701 | 11.871 | 2.398 | 1.008 | 0.530 | 0.112 | 2.184 | 10.812 | Astrocytes | Astrocyte |
| **KCNS2** | 8 | 0.100 | 2.302 | 0.105 | 0.100 | 0.100 | 0.100 | 0.100 | 21.829 | 23.022 | Neuron | |
| **LAIR2** | 19 | | | | | | | | | | | |
| **LCP2** | 5 | 0.102 | 0.323 | 2.880 | 0.139 | 0.321 | 52.317 | 14.336 | 3.649 | 18.164 | Microglia | Microglia |
| **LY6H** | 8 | 1.736 | 225.441 | 72.095 | 17.532 | 7.308 | 8.442 | 2.675 | 3.127 | 12.859 | Neuron | |
| **MET** | 7 | 0.196 | 1.235 | 0.229 | 0.143 | 0.100 | 0.100 | 0.346 | 3.568 | 5.397 | Neuron | |
| **MGP** | 12 | 0.428 | 0.777 | 2.918 | 0.397 | 0.882 | 0.668 | 30.488 | 10.449 | 34.549 | Endothelial Cells | |
| **SERPINF1** | 17 | 9.423 | 2.487 | 10.790 | 0.539 | 1.386 | 97.531 | 7.738 | 9.039 | 10.350 | Microglia | |
| **PRRX1** | 1 | 17.844 | 0.918 | 15.575 | 2.600 | 0.111 | 0.100 | 0.556 | 1.146 | 6.863 | | |
| **ROBO1** | 3 | 2.399 | 11.005 | 11.671 | 2.815 | 0.293 | 0.100 | 0.432 | 1.061 | 4.145 | Neuron | Oligodendrocyte Precursor Cell |
| **TGFBI** | 5 | 0.446 | 0.492 | 2.170 | 1.386 | 2.077 | 24.550 | 0.845 | 11.312 | 11.819 | Microglia | Microglia |
| **TRPC6** | 11 | 0.100 | 0.677 | 0.138 | 0.100 | 0.100 | 0.100 | 0.100 | 4.890 | 6.768 | Neuron | |
| **ZYX** | 7 | 12.443 | 12.732 | 17.570 | 12.735 | 20.805 | 74.186 | 35.417 | 2.095 | 3.566 | Microglia | |
| **DIRAS3** | 1 | | | | | | | | | | | |
| **FAM65B** | 6 | 6.286 | 30.186 | 1.074 | 0.105 | 0.276 | 0.976 | 0.869 | 4.802 | 28.112 | Neuron | |
| **ARHGAP25** | 2 | 0.187 | 0.230 | 2.900 | 0.136 | 0.362 | 76.103 | 18.077 | 4.210 | 26.240 | Microglia | |
| **FRAT1** | 10 | 1.646 | 5.892 | 1.594 | 0.663 | 1.206 | 1.676 | 1.014 | 3.516 | 3.579 | Neuron | |
| **SPON2** | 4 | 0.116 | 0.367 | 0.134 | 0.214 | 0.100 | 0.104 | 1.592 | 4.336 | 7.442 | Neuron | |
| **CNTN6** | 3 | 1.022 | 2.394 | 13.467 | 4.841 | 0.100 | 0.137 | 0.100 | 2.782 | 5.626 | Oligodendrocyte Precursor Cell | |
| **CDR2L** | 17 | 2.975 | 10.019 | 16.521 | 17.275 | 15.453 | 0.516 | 3.515 | 1.046 | 1.118 | | Oligodendrocyte |
| **VILL** | 3 | 0.100 | 0.100 | 0.100 | 0.100 | 0.100 | 0.100 | 0.100 | 1.000 | 1.000 | | |
| **HR** | 8 | 5.561 | 1.842 | 7.492 | 24.163 | 12.185 | 1.056 | 0.100 | 1.983 | 3.225 | Newly Formed Oligodendrocyte | |
| **NT5M** | 17 | 11.875 | 16.775 | 16.512 | 13.975 | 15.025 | 11.235 | 11.594 | 1.016 | 1.116 | | |

| | | | | | | | | | | | | |
|---|---|---|---|---|---|---|---|---|---|---|---|---|
| **SHD** | 19 | 9.325 | 14.378 | 19.730 | 9.362 | 3.409 | 0.886 | 0.910 | 1.372 | 2.107 | | Interneuron |
| **ELMO3** | 16 | 0.623 | 0.173 | 0.183 | 0.100 | 0.100 | 0.164 | 0.330 | 1.886 | 3.409 | Astrocytes | |
| **MAGED4B** | X | | | | | | | | | | | |
| **C12ORF23** | 12 | | | | | | | | | | | |
| **GLCCI1** | 7 | 4.252 | 4.604 | 7.851 | 5.422 | 1.839 | 0.340 | 7.182 | 1.093 | 1.448 | | |
| **SLC22A9** | 11 | | | | | | | | | | | |
| **GPRIN1** | 5 | 0.588 | 28.585 | 4.909 | 1.203 | 1.539 | 1.280 | 0.280 | 5.823 | 18.574 | Neuron | |
| **LOC145837** | 15 | | | | | | | | | | | |
| **CBLN2** | 18 | 0.101 | 11.801 | 0.354 | 4.057 | 0.508 | 0.185 | 2.503 | 2.909 | 4.716 | Neuron | |
| **FAM162B** | 6 | 0.146 | 0.103 | 0.100 | 0.100 | 0.100 | 0.100 | 0.351 | 2.400 | 3.402 | Endothelial Cells | |
| **PGM2L1** | 11 | 1.130 | 14.572 | 2.629 | 0.567 | 1.142 | 0.742 | 1.982 | 5.542 | 7.351 | Neuron | |
| **IGSF22** | 11 | | | | | | | | | | | |
| **KIAA1875** | 8 | | | | | | | | | | | |
| **CTXN3** | 5 | 0.100 | 0.307 | 0.137 | 0.102 | 0.199 | 0.100 | 0.100 | 1.544 | 2.240 | Neuron | |
| **IQCJ** | 3 | 0.100 | 0.101 | 0.346 | 0.100 | 0.100 | 0.100 | 0.100 | 3.415 | 3.462 | Oligodendrocyte Precursor Cell | |

Database tables: Online supplemental material of Zhang et al., 2014 (http://web.stanford.edu/group/barres_lab/brain_rnaseq.html; based on tissues from the mouse cerebral cortex); Table S1 of Zeisel et al., 2015 (based on tissues from the mouse somatosensory cortex and hippocampal CA1 region).

**Table S9. Cell-type specificity in expression of the consensus set of genes in the human brain neocortex.**

| GENES | ZHONG2018 | LI2018 | FAN2018 | LAKE2018 |
|---|---|---|---|---|
| **ANK1** | | | | **In3/Inba/In6b** |
| **COX7A1** | | | **Cajal_Retzius**/Endo_1 | |
| **LAIR2** | **Excitatory neurons_WEEK** | | Effector T cell | |
| **LCP2** | | Microglia_prenatal | | |
| **LY6H** | Microglia | | **Cajal_Retzius** | |
| **MGP** | | VSMC_adult | | |
| **SERPINF1** | Microglia | Microglia_prenatal | Microglia | |
| **ROBO1** | **NPC(neural progenitor cells)/Excitatory neurons_WEEK/Interneurons** | **IntN_adult** | | **In1a/In1b/In1c/In2/In3/In4a/In7/In8/**OPC_Cer |
| **TGFBI** | | | Myeloid cell | |
| **TRPC6** | | | Endo_2 | |
| **ZYX** | Astrocytes | | Endo_2 | |
| **FAM65B** | Microglia | | Immune/B cell | |
| **ARHGAP25** | | Microglia_prenatal | | |
| **SPON2** | | | Endo_2 | |
| **CNTN6** | | | | **In7** |
| **CDR2L** | | | Olig | |
| **HR** | | | Olig | |
| **GLCCI1** | Astrocytes | | | **Ex3b/Ex3d/Ex4/**OPC_Cer |
| **GPRIN1** | Microglia | | | |
| **CBLN2** | **Excitatory neurons_stage**/Microglia | | | **Ex1/Ex2/Ex8** |
| **FAM162B** | | | Endo_2 | |
| **PGM2L1** | **Excitatory neurons_WEEK**/Astrocytes/Microglia | | | **Ex3e/Ex5a/Ex6a/Ex6b/In1b/In4b** |

Database tables: Table S2 of Zhong et al., 2018 (based on tissues from prefrontal regions); Table S8 from Li et al., 2018 (various cortical areas); Table S3 from Fan et al., 2018 (various cortical areas); Table S3 from Lake et al., 2018 (frontal and visual cortex, and cerebellum).

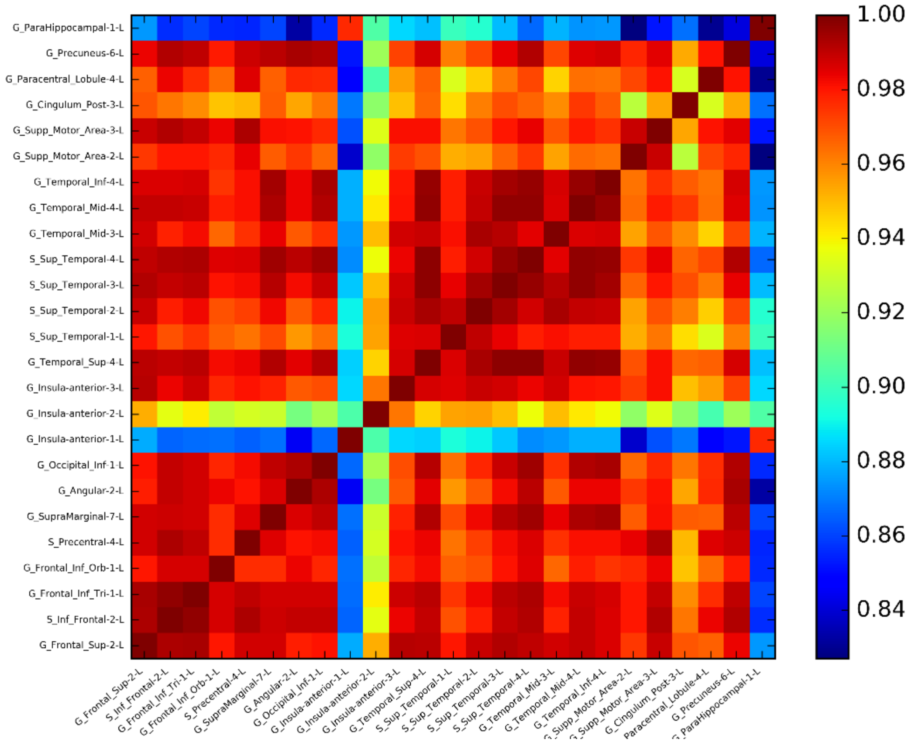


**Fig. S1. Two deep cortical areas (G_ParaHippocampal-1 and G_Insula-anterior-1) showing relatively distinct gene expression profiles compared with most of the cortical areas.** The gene expression similarity was calculated based on the top 5% genes according to their differential stability across donors.

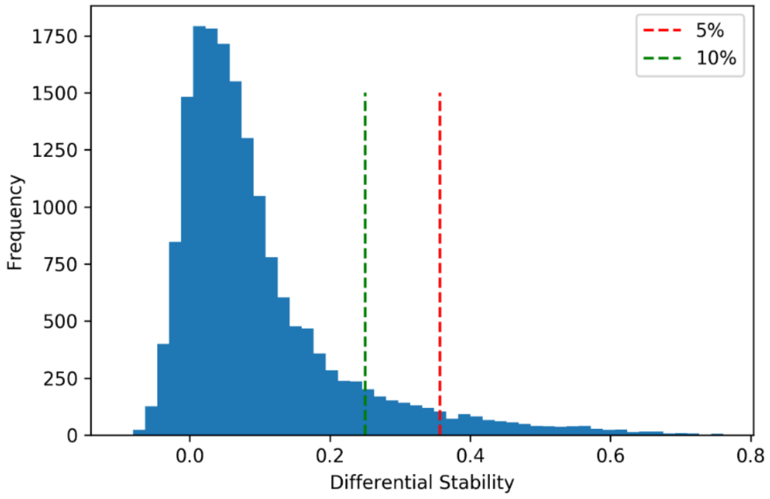


**Fig. S2. Distribution of the differential stability score (across the cerebral cortex structures) for all genes in the Allen Human Brain Atlas.** The thresholds of top 5% and 10% are indicated with red and green lines, respectively.

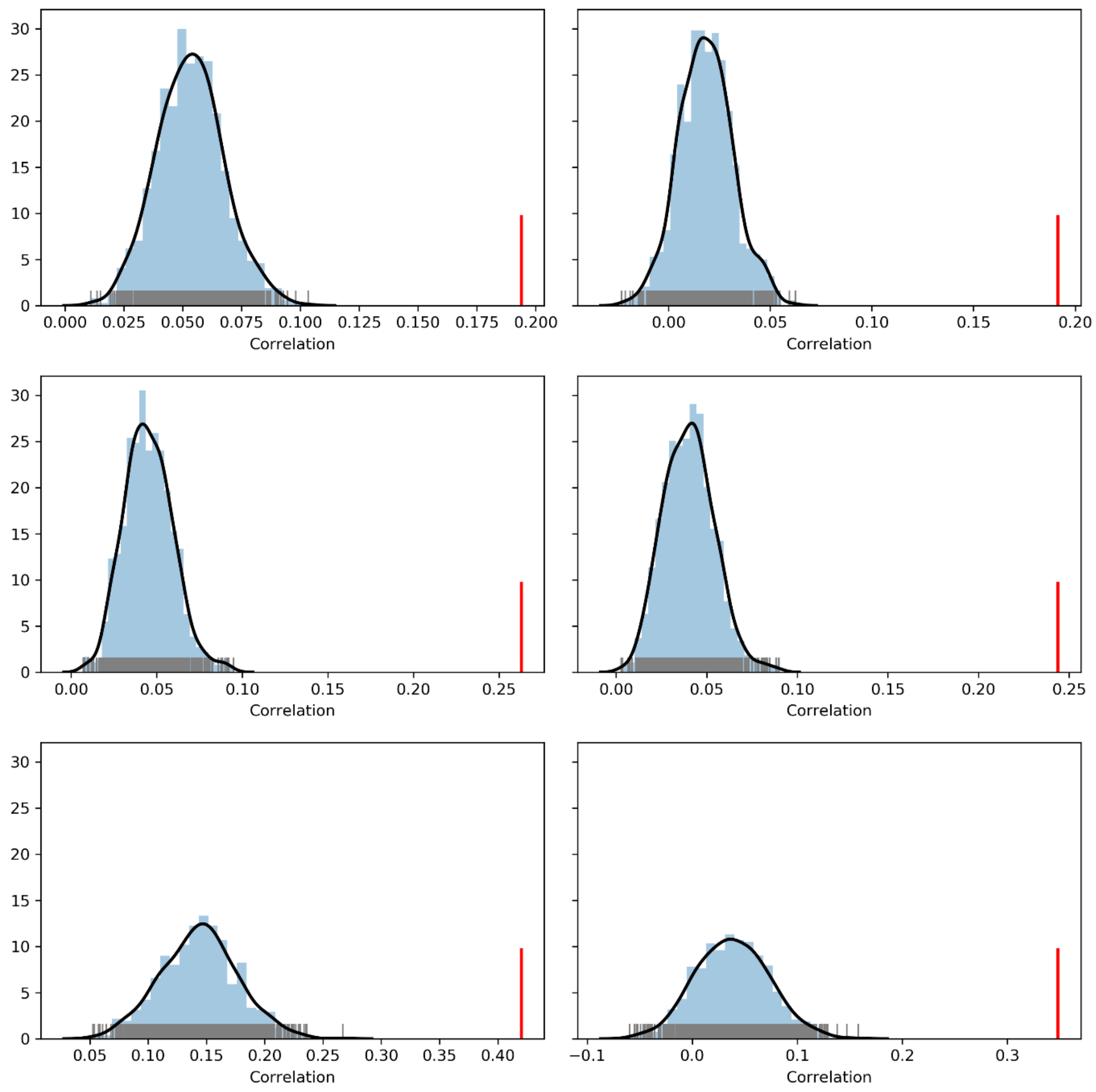


**Fig. S3.** Distributions of the connectivity-transcriptome correlations based on repeat random sampling of the same number of genes as in the top 5% list (based on differential stability across donors). The red lines indicate the correlations based on the top 5%. The figure is organized by language network definitions (rows) and resting-state fMRI datasets (columns).

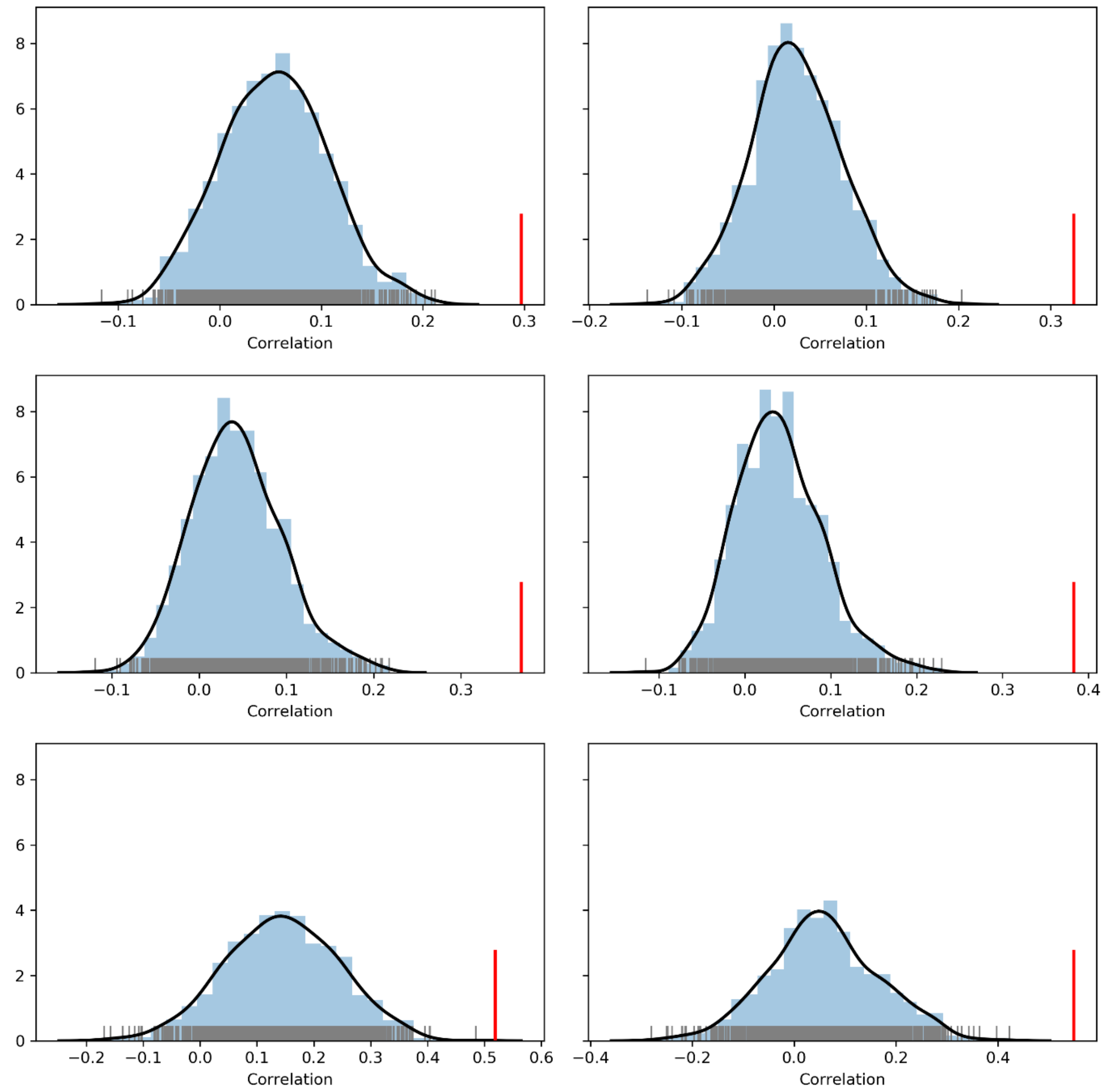


**Fig. S4.** Distributions of the overall connectivity-transcriptome correlations based on repeat random sampling of the same number of genes as in the consensus gene set (41 genes). The red lines indicate the correlations based on the consensus set. The figure is organized by language network definitions (rows) and resting-state fMRI datasets (columns).
